# Determinants of RNA recognition by the FinO domain of the *Escherichia coli* ProQ protein

**DOI:** 10.1101/2020.05.04.075150

**Authors:** Ewa M. Stein, Joanna Kwiatkowska, Maciej M. Basczok, Chandra M. Gravel, Katherine E. Berry, Mikołaj Olejniczak

## Abstract

The regulation of gene expression by small RNAs in *Escherichia coli* depends on RNA binding proteins Hfq and ProQ, which bind mostly distinct RNA pools. To understand how ProQ discriminates between RNA substrates, we compared its binding to six different RNA molecules. Full-length ProQ bound all six RNAs similarly, while the isolated N-terminal FinO domain (NTD) of ProQ specifically recognized RNAs with Rho-independent terminators. Analysis of *malM* 3’-UTR mutants showed that tight RNA binding by the ProQ NTD required a terminator hairpin of at least two base pairs preceding an 3’ oligoU tail of at least four uridine residues. Substitution of an A-rich sequence on the 5’ side of the terminator to uridines strengthened the binding of several ProQ-specific RNAs to the Hfq protein, but not to the ProQ NTD. Substitution of the motif in the *malM*-3’ and *cspE*-3’ RNAs also conferred the ability to bind Hfq in *E. coli* cells, as measured using a three-hybrid assay. In summary, these data suggest that the ProQ NTD specifically recognizes 3’ intrinsic terminators of RNA substrates, and that the discrimination between RNA ligands by *E. coli* ProQ and Hfq depends both on positive determinants for binding to ProQ and negative determinants against binding to Hfq.

## INTRODUCTION

RNA-binding proteins are important contributors to major life processes, including the regulation of gene expression by RNAs (1). In many bacteria, the prominent role played by *trans*-encoded base-pairing small RNAs (sRNAs) in regulating gene expression requires a matchmaker protein called Hfq, which assists sRNA in pairing to complementary sequences in target mRNAs and affects sRNA stability (2-6). Global identification of new RNA/protein interactions has been enabled by deep-sequencing approaches (7), such as Grad-seq, which uses glycerol gradients to identify novel RNA/protein complexes (8), CLIP-seq, which uses crosslinking to define protein binding sites in the transcriptome (9), and RIL-seq which identifies protein-dependent RNA-RNA interactions (10). Recent studies using these approaches showed that another protein, named ProQ, is a global RNA binding protein in *Escherichia coli* and *Salmonella enterica* (11-13), and is involved in sRNA interactions with other RNA molecules (12,14,15). The pool of RNA ligands of ProQ is mostly distinct from that of Hfq and contains more mRNAs than sRNAs (11,12).

While ProQ was originally discovered in a search for genes that affect proline transport (16), this protein contributes to several physiological processes in *E*.*coli* and *S. enterica* (17) – including DNA metabolism (13,14), bacterial virulence (15), and adaptation to osmotic stress (12) and resource limitation (18). ProQ is active as a monomer (19), and it belongs to the FinO-domain family with representatives in numerous γ-proteobacteria (17,20). Other members of this family include the F-like plasmid FinO protein (21,22), *Legionella pneumophila* RocC protein (23), and *S. enterica* pCol1B9 plasmid-encoded FopA protein (J. Vogel, personal communication), which each interact with few RNAs to control expression of specific genes, and a minimal ProQ protein (NMB1681), which, like *E. coli* ProQ, is a global RNA-binding protein in *Neisseria meningitidis* (24,25).

The *E. coli* ProQ protein consists of the N-terminal FinO domain (NTD) (19,26,27), for which this protein family is named (28), and the C-terminal Tudor domain (CTD), connected by an extended linker enriched for positively charged residues (19). The surface of the FinO domain of ProQ displays patches of positively and negatively charged residues, which could contribute to RNA binding via electrostatic interactions or hydrogen bonding (17). It was shown that the isolated FinO domain binds a double-stranded model RNA as well as the full-length *E. coli* ProQ protein does, suggesting that the NTD is the main site of RNA binding in ProQ (29). In agreement with that suggestion mapping of the ProQ surfaces affected by the binding of two natural ProQ ligands, the 3’-UTR of *cspE* mRNA and the sRNA SraB, using hydrogen-deuterium exchange (HDX) experiments, showed the strongest protection by each RNA on partly overlapping surfaces of the FinO domain (19). In further support, a study using a bacterial three-hybrid assay showed that mutations in the FinO domain of ProQ were detrimental for the binding of the *cspE* 3’-UTR and SibB sRNA in *E. coli* cells, and that this domain alone was sufficient for the binding of each RNA (30).

Additionally, mutations in the homologous domain of *L. pneumophila* RocC negatively affected the RocR sRNA lifetime in cells (23), and the homologous domain of the FinO protein crosslinked to bound RNA (31). While the FinO domain of ProQ appears to be the main site of RNA binding, reported data suggest that other regions of ProQ also form interactions with RNA molecules. The HDX study suggested that the sRNA SraB contacts the linker and CTD of *E. coli* ProQ more extensively than the *cspE* 3’-UTR (19). In agreement with this observation, the linker and the CTD regions of ProQ contributed to the binding of SibB sRNA, but not the *cspE* 3’-UTR, when measured by the three-hybrid assay (30).

The ProQ protein binds structured RNA motifs in its ligands in *E. coli* and *S. enterica* (11,12). This preference for double-stranded RNA was initially observed in studies of *E. coli* ProQ binding to RNAs derived from FinP RNA (29). RNA ligands of *S. enterica* ProQ identified by the Grad-seq method were also observed to be highly structured (13). Further analysis of ProQ binding sites in the transcriptomes of *E. coli* and *S. enterica* using CLIP-seq method showed that ProQ bound to GC-rich double-stranded elements, including Rho-independent terminators (11). Similarly, RIL-seq analysis of transcripts crosslinked to *E. coli* ProQ showed that interacting regions of RNAs contained GC-rich motifs followed by uridine tracts (12). Other FinO-domain proteins – minimal *N. meningitidis* ProQ, *L. pneumophila* RocC, and F-like plasmid FinO – also preferably bind secondary structure motifs, including transcription terminators (23,25,32).

In contrast to the behavior of ProQ, the Hfq protein binds single-stranded RNA sequences, which include uridine- or adenosine-rich motifs (33,34). Hfq often binds single-stranded A-rich sequences, including ARN repeats, in 5’-UTRs of mRNA molecules and in ribosomal RNAs (35-37). On the other hand, the binding of Hfq to sRNAs is dependent on the 3’ terminal oligoU tails (38), and either single-stranded U-rich sequences (39), or single-stranded A-rich sequences (40-42) upstream of the terminator hairpin. Hence, although Hfq binds at Rho-independent transcription terminators in some of its RNA ligands, Hfq binding to these regions involves single-stranded sequences.

ProQ and Hfq bind to different, partly overlapping, pools of RNA ligands in *E. coli* and *S. enterica* (11-13). It was observed that the group of highly structured ProQ-binding RNAs identified using Grad-seq was distinct from that of Hfq-binding RNAs (13). Further studies using CLIP-seq revealed that ProQ recognizes its RNA ligands using a structural code, which is different from Hfq’s recognition of RNA sequence motifs (11). Additionally, the RIL-seq analysis of RNA-RNA interactions on ProQ and Hfq showed that while the RNA sets most enriched on each protein were generally unique, there was some overlap (12). The distinctive features of ProQ RNA ligands include structured GC-rich RNA motifs and single-stranded oligoU tails of intrinsic terminators that are shorter than those in Hfq-specific RNAs (11,12). As it has been proposed that some of ProQ functions center on RNA 3’ ends (11), it is likely that properties of 3’ RNA regions could be important for the discrimination between RNA ligands by ProQ and Hfq proteins.

To better understand how *E. coli* ProQ recognizes its RNA ligands, we compared the binding of six natural RNA ligands to the full-length ProQ and the isolated NTD domain using an electrophoretic mobility shift assay. The data showed that the N-terminal FinO domain is the part of ProQ which is responsible for the recognition of Rho-independent terminator structures in RNA ligands of ProQ. Studying mutations in the Rho-independent terminator of *malM* 3’-UTR showed that binding of the NTD required a double-stranded region of at least two base-pairs in length, and a 3’ oligoU tail of four or more uridine residues. Finally, results obtained using *in vitro* and *in vivo* binding assays showed that an A-rich sequence located on the 5’ side of the terminator hairpin in several ProQ-specific RNAs serves as a negative determinant of their binding to the Hfq protein. Overall, these data suggest that the sequences and structures of ProQ-specific RNAs have been optimized to ensure correct recognition by ProQ, while at the same time preventing their binding by Hfq.

## MATERIALS AND METHODS

### Preparation of RNAs

The DNA templates for the *in vitro* transcription were obtained by Taq polymerase extension of chemically synthesized overlapping oligodeoxyribonucleotides (Sigma-Aldrich) (Supplemental Table S1). RNAs were transcribed with T7 RNA polymerase and purified using denaturing gel electrophoresis, as described (43,44). RNA molecules were 5’-^32^P labeled using T4 polynucleotide kinase (Thermo Scientific), which was followed by phenol-chloroform extraction, denaturing gel purification, and ethanol precipitation.

### Protein overexpression and purification

The sequences of *E. coli* ProQ protein or its N-terminal domain (NTD) were cloned into pET15b vector (Novagen) using BamHI restriction site (Supplemental Table S2). In the constructs the coding sequence of the protein was preceded by His_6_-tag and TEV protease recognition sequence (ENLYFQ↓S). Constructs were overexpressed in BL21 *Δhfq E. coli* strain (a kind gift of Prof. Agnieszka Szalewska-Pałasz, University of Gdańsk). After harvesting, the cells were suspended in buffer consisting of 50 mM Tris, pH 7.5, 500 mM NaCl and 10% glycerol (buffer A), and frozen at −80°C. Thawed cells were mixed with protease inhibitor cocktail (Roche) and lysed by sonication, followed by clarification of the lysates by centrifugation. After lysis, the freezing and thawing of the samples was avoided during all steps in the purification procedure. Lysates were loaded onto HisTrap crude column (GE Healthcare) on FPLC system (AKTA pure), and the proteins were eluted with linear imidazole gradient (25-600 mM) in buffer A. To remove nucleic acids, the samples were loaded onto HiTrap heparin column (GE Healthcare), and eluted with linear NaCl gradient (0.1-2 M). Subsequently, the His_6_-tag was removed by TEV-His_6_ protease used at 100:1 OD ratio of total protein to TEV, at 4°C, overnight. The completion of the TEV cleavage was confirmed by SDS-PAGE, and the samples were loaded onto HisTrap column and eluted in buffer A containing 25 mM imidazole. The preparation was further purified using gel filtration on HiLoad 16/60 Superdex 200 size exclusion column (GE Healthcare) and eluted in storage buffer containing 50 mM Tris, pH 7.5, 300 mM NaCl, 10% glycerol and 1 mM EDTA. The samples were stored at −80 °C as 10 µl and 20 µl aliquots, and used without refreezing. The molecular weights of the purified proteins were determined by MALDI-TOF as 25964.9 Da for ProQ, and 14673.5 Da for NTD, which agrees well with the calculated mass of 25979.5 Da for ProQ, and 14669.5 Da for the NTD, each with an additional N-terminal serine residue remaining from TEV cleavage site. The protein concentration was determined by measuring the absorption at 280 nm using extinction coefficient of 9650 M^-1^ cm^-1^.

The *E. coli* Hfq protein was purified as previously described (40).

### ProQ-binding assay

Prior to use, RNAs were denatured at 90°C for 2 min followed by refolding on ice for 5 min. The concentration series of ProQ or NTD were made by 2-fold dilutions from the highest concentration. 1 nM ^32^P-labeled RNA was incubated with indicated concentrations of ProQ in the binding buffer (25 mM Tris, pH 7.5, 150 mM NaCl, 5% glycerol, 1 mM MgCl_2_) for 30 min at RT. Incubations were performed in low-protein binding microplates, additionally pre-treated with a solution containing 0.0025% BSA. After incubation, the samples were loaded onto a native 6% polyacrylamide gel (19:1), and run in 0.5×TBE at 4°C. Gels were vacuum-dried, exposed to phosphor screens and quantified using a phosphorimager with MultiGauge software (Fuji FLA-5000). In the data fitting the *GraphPad Prism* software was used. The equilibrium dissociation constant (*K*_d_) values were calculated by fitting data to the quadratic equation (45),

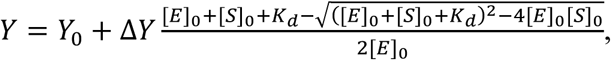

where *Y* is a fraction bound, *Y*_*0*_ is a bottom plateau, and Δ*Y* is the difference between top and bottom plateaus. [*E*]_0_ is concentration of ^32^P-labeled RNA, and [*S*]_0_ is the concentration of the ProQ protein or the NTD.

### Hfq-binding assay

RNA binding to the *E. coli* Hfq protein was measured using double-filter retention assay, essentially as previously described (40,44). Prior to use ^32^P-RNA was denatured by heating at 90°C for 2 min followed by refolding on ice for 5 min. Then 20 μL of 200 pM ^32^P-RNA was mixed with 20 μL of the Hfq dilutions in binding buffer (25 mM Tris-HCl pH 7.5, 150 mM NaCl, 5% glycerol, 1 mM MgCl_2_) and incubated for 30 min at RT in low-protein binding microplates, additionally pre-treated with a solution containing 0.0025% BSA. Afterwards, 35 μl aliquots were filtered and washed with 100 μL of 1× binding buffer. Membranes were dried and exposed to phosphorscreens, followed by quantification using a phosphorimager (Fujifilm FLA-5000) with ImageQuant software. The *K*_d_ values were determined by fitting data to the quadratic equation as described above for ProQ and the NTD.

### Computational analysis of sequences of ProQ- and Hfq-specific RNAs

To analyze the nucleotide content of the sequence immediately 5’-adjacent to the terminator hairpin we used the databases of ProQ and Hfq binding sites in the transcriptomes of *E. coli* and *S. enterica* obtained using RIL-seq and CLIP-seq methods. The sequences of *E. coli* RNAs that bind to ProQ were obtained from published RIL-seq data (12) and CLIP-seq data (11). The sequences of *E. coli* sRNAs binding to Hfq were obtained from RIL-seq data (12). The sequences of *S. enterica* sRNAs binding to ProQ and those binding to Hfq were obtained from CLIP-seq data (9,11). For the analysis we selected the top 50 RNA ligands of each protein containing Rho-independent terminators (Suppl. Table S3). Specifically, from the database of individual RNAs from RIL-seq in LB (12), the top 50 RNA sequences annotated as mRNA 3’ UTRs or sRNAs were chosen for the analysis. From CLIP-seq data (9,11) 50 RNA sequences with the highest averaged read-counts were selected, in which the CLIP-seq peak covered a terminator loop or when it was located no further than 10 nucleotides upstream from it. We note that, of the top 50 ProQ RNA ligands with intrinsic terminators, most were mRNA 3’ UTRs, while of the top 50 Hfq RNA ligands with terminators, most were sRNAs.

In these RNA ligands, a 10-nt sequence 5’-adjacent to the intrinsic terminator was analyzed. To define the sequences 5’-adjacent to the terminator hairpins the secondary structure of each RNA was analyzed using the *RNAStructure* software (46). The 3’ end of this sequence was defined as the nucleotide which was located opposite to the first uridine of the 3’ oligoU tail on the other side of the terminator hairpin. Afterwards, the 10-nt sequences 5’-adjacent to the terminator hairpins were extracted and analyzed using *WebLogo* software (47) for the five datasets: RIL-seq data for ProQ and for Hfq in *E. coli* (12), CLIP-seq data for ProQ in *E. coli* (11), and CLIP-seq data for ProQ and for Hfq in *S. enterica* (9,11). Nucleotide frequency was calculated for each dataset using *WebLogo* software and compared with the whole-transcriptome nucleotide frequency in χ^2^ goodness-of-fit test. Subsequently, the statistical importance of nucleotide frequency at each position of the 10-nt sequence 5’-adjacent to the terminator hairpin in ProQ ligands was calculated in reference to each position of this sequence in Hfq ligands using the *pLogo* software (48). The *pLogo* analysis was performed for the *E. coli* ProQ dataset versus Hfq dataset obtained by RIL-seq (12) and for the *S. enterica* ProQ dataset versus Hfq dataset obtained by CLIP-seq (9,11).

### Bacterial strains and plasmids used in the bacterial three hybrid assay

A complete list of plasmids, strains, and oligonucleotides used in this assay is provided in Supplemental Tables S4-S7. NEB 5-alpha F’Iq cells (New England Biolabs) were used as the recipient strain for all plasmid constructions. Reporter strains KB483 (O_L_2-62-*lacZ*; Δ*hfq*) and KB480 (O_L_2-62-*lacZ*; *hfq*^+^) were used as the reporter strains for bacterial three-hybrid (B3H) experiments (30) (Suppl. Table S4). Plasmids were constructed as specified in the Supplemental Tables S5 and S6. The *E. coli (Ec) malM* 3’-UTR (final 90 nts) was PCR amplified using *Ec* genomic DNA and ligated into to pCDF-1XMS2^hp^ (pCH1) between XmaI and HindIII to generate MS2^hp^-*malM* 3’-UTR (pKB1210). PCR mutagenesis to create site-directed mutants of *malM* 3’-UTR and *cspE* 3’-UTR was conducted with the Q5 site-directed mutagenesis kit (New England Biolabs) using end-to-end primers designed with NEBaseChanger. The full sequence of each hybrid RNA is provided in Supplementary Table S7.

### Bacterial three-hybrid assay in *E. coli*

For B3H assays, reporter cells (KB480 or KB483) were freshly co-transformed with compatible pAC-, pBR- and pCDF-derived plasmids, as indicated. Transformed bacteria were inoculated into 1 ml LB broth supplemented with carbenicillin (100 µg/ml), chloramphenicol (25 µg/ml), tetracycline (10 µg/ml), spectinomycin (100 µg/ml) and 0.2% arabinose in a 2 ml 96-well deep well block (VWR), sealed with breathable film (VWR) and shaken at 900 rpm at 37°C. Overnight cultures were diluted 1:50 into 200 µl LB supplemented as above and cells were grown to mid-log (OD600 ≈ 0.6) in optically clear 200 µl flat bottom 96-well plates (Olympus) covered with plastic lids, as above. From here, 2 µl of cells were diluted 1:100 and plated on 5-bromo-4-chloro-3-indolyl-β**-**D-galactopyranoside (X-Gal)-indicator medium (containing arabinose and antibiotics as above along with 40 µg/mL X-Gal, 70 µM phenylethyl-β-D-thiogalactopyranoside (TPEG), and 1.5 µM Isopropyl β-D-1-thiogalactopyranoside (IPTG)) and incubated overnight. Following overnight growth, plates were allowed to sit at 4°C for 1-2 days to allow color to develop, and were photographed with a black-velvet background and oblique lighting. Brightness and levels were adjusted evenly across photographs. B3H interactions are interpreted as the β-gal activity in reporter cells containing all hybrid constructs (α-ProQ, λCI-MS2^CP^ and MS2^hp^-bait hybrid RNA), relative to the highest activity from negative controls – cells containing plasmids where one of the hybrid constructs is replaced by an α empty, CI empty or MS2^hp^ empty construct. In the context of qualitative plate-based assays, this is represented as the relative color of bacterial patches on X-gal containing media, where blue color represents higher β-gal activity, and therefore stronger RNA-protein interaction. Assays were conducted in duplicate on at least three separate days and a representative experiment is shown.

## RESULTS

To study the recognition of RNA ligands by the *E. coli* ProQ protein, 6 functionally and structurally diverse RNA molecules were selected (Fig. 1). In this set two were sRNAs expressed as independent transcripts – MicA and SibA, two were mRNA 3’-UTRs – *malM*-3’ and *cspE*-3’, and two were mRNA 5’-UTRs – *lpp*-5’ and *hupA*-5’. These RNAs have been identified in *E. coli* as ProQ ligands using CLIP-seq (11) and detected as single fragments or in chimeras on ProQ using RIL-seq (12). Among them, MicA is a *trans*-encoded base pairing sRNA, which is an important ligand of the Hfq protein (49), while SibA is a *cis*-encoded base pairing sRNA (50). *malM*-3’ is a processing product of 3’-UTR of the respective mRNA, and *cspE*-3’ is made as an independent transcription unit within 3’-UTR (12). The 5’-UTR of *lpp* mRNA was found in two of the most abundant chimeras identified on ProQ by RIL-seq (12). Additionally, the homologous region of the *hupA* 5’-UTR was identified in *S. enterica* as a target for regulation by the ProQ-dependent sRNA RaiZ (14). Hence, beyond their different sequences and structures, these RNAs also differ in their origins, making them representative of a diversity of ProQ RNA ligands.

**Figure 1.**
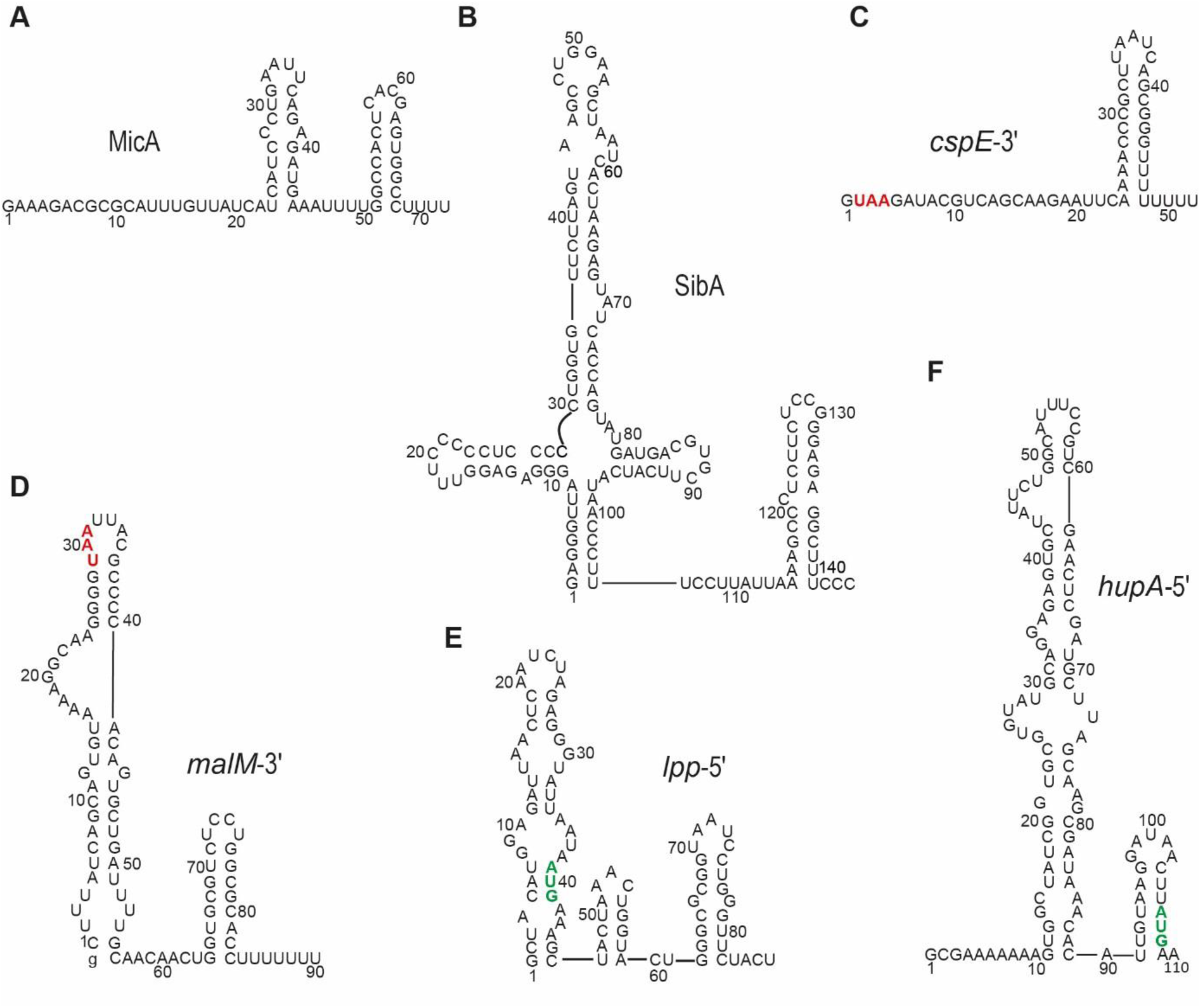
RNA molecules bound by ProQ protein in *E. coli*, which were used in this study. The secondary structures of small RNAs (A) MicA (49), and (B) SibA (13) are presented according to the references. The secondary structures of (C) *cspE*-3′, (D) *malM*-3′, (E) *lpp-*5’, and (F) *hupA*-5’ were predicted using *RNAstructure* software (46). UAA stop codons and AUG start codons are shown in red and green, respectively. The lower case g denotes guanosine residue added on 5′ end of *malM*-3′ to enable T7 RNA polymerase transcription.

### ProQ binds diverse RNA molecules with similar affinities

The binding affinity of the full-length *E. coli* ProQ protein to RNA molecules was determined using a gel shift assay (Fig. 2). The electrophoretic mobility shift data showed that ProQ complexes with each of the RNAs were well resolved from unbound RNAs on the gel. In the range of ProQ concentrations used, five RNAs: MicA (72-nt), SibA (144-nt), *cspE*-3’ (52-nt), *malM*-3’ (91-nt), and *lpp*-5’ (86-nt), each formed one dominant complex with ProQ, while the 110-nt long *hupA*-5’ formed an additional higher order complex with ProQ (Fig. 2). The equilibrium binding constant (*K*_d_) values ranged between 2.4 nM (for SibA) and 21 nM (for *lpp*-5’) (Fig. 2, Table 1). Aside from SibA, which bound ProQ more than 8-fold tighter than *lpp*-5’, the binding affinities of other RNAs with ProQ differed less than 3-fold among one another and were similar to the previously reported *K*_d_ of 31 nM for *E. coli* ProQ binding to a fragment of FinP RNA (29), and to low nanomolar *K*_d_ values estimated for SibA and RaiZ binding to *S. enterica* ProQ in a buffer with lower ionic strength (14). Overall, these results showed that the full-length *E. coli* ProQ binds these six natural RNA ligands with similar affinities.

**Table 1.** Equilibrium binding of different RNAs and their mutants with extended or mutated 3′-ends to the full-length ProQ and its N-terminal domain (NTD).

| <sup>32</sup>P-RNA | $K_d$ [nM] | |
| --- | --- | --- |
|  | ProQ | NTD |
| MicA | $15 \pm 5.1$ | $21 \pm 10$ |
| SibA | $2.4 \pm 0.85$ | $4.7 \pm 0.89$ |
| *cspE*-3' | $13 \pm 4.5$ | $4.5 \pm 1.5^a$ |
| *malM*-3' | $12 \pm 5.3^a$ | $12 \pm 4.8$ |
| *lpp*-5' | $21 \pm 5.9^a$ | $> 1000$ |
| *hupA*-5' | $9.3 \pm 4.1^b$ | $> 1000$ |
| *lpp*-5'-mut | $3 \pm 1.3$ | $1.3 \pm 0.83$ |
| *lpp*-5'-term | $3.6 \pm 1.3$ | $5.1 \pm 1$ |
| *lpp*-5'-hp | $20 \pm 2.8^b$ | n. d. |
| *hupA*-5'-mut | $14 \pm 8^b$ | $16 \pm 4.7$ |
| *hupA*-5'-term | $10 \pm 5$ | $8.5 \pm 0.96$ |
| *hupA*-5'-hp | $18 \pm 1.9^b$ | n. d. |
| *cspE*-3'-ext+5 | $31 \pm 14^a$ | $17 \pm 7.5$ |
| *cspE*-3'-ext+22 | $17 \pm 8.7$ | n. d. |
| *malM*-3'-ext+5 | $33 \pm 4.2$ | n. d. |
| *malM*-3'-ext+17 | $8.4 \pm 2.6^b$ | n. d. |

<sup>a</sup> – The endpoint value of 100% was assumed in the data fitting.
<sup>b</sup> – In the $K_d$ value calculations the fractions of both RNA-protein complexes were summed together.
n. d. – the binding was essentially undetectable up to 1 μM concentration of the NTD.

**Figure 2.**
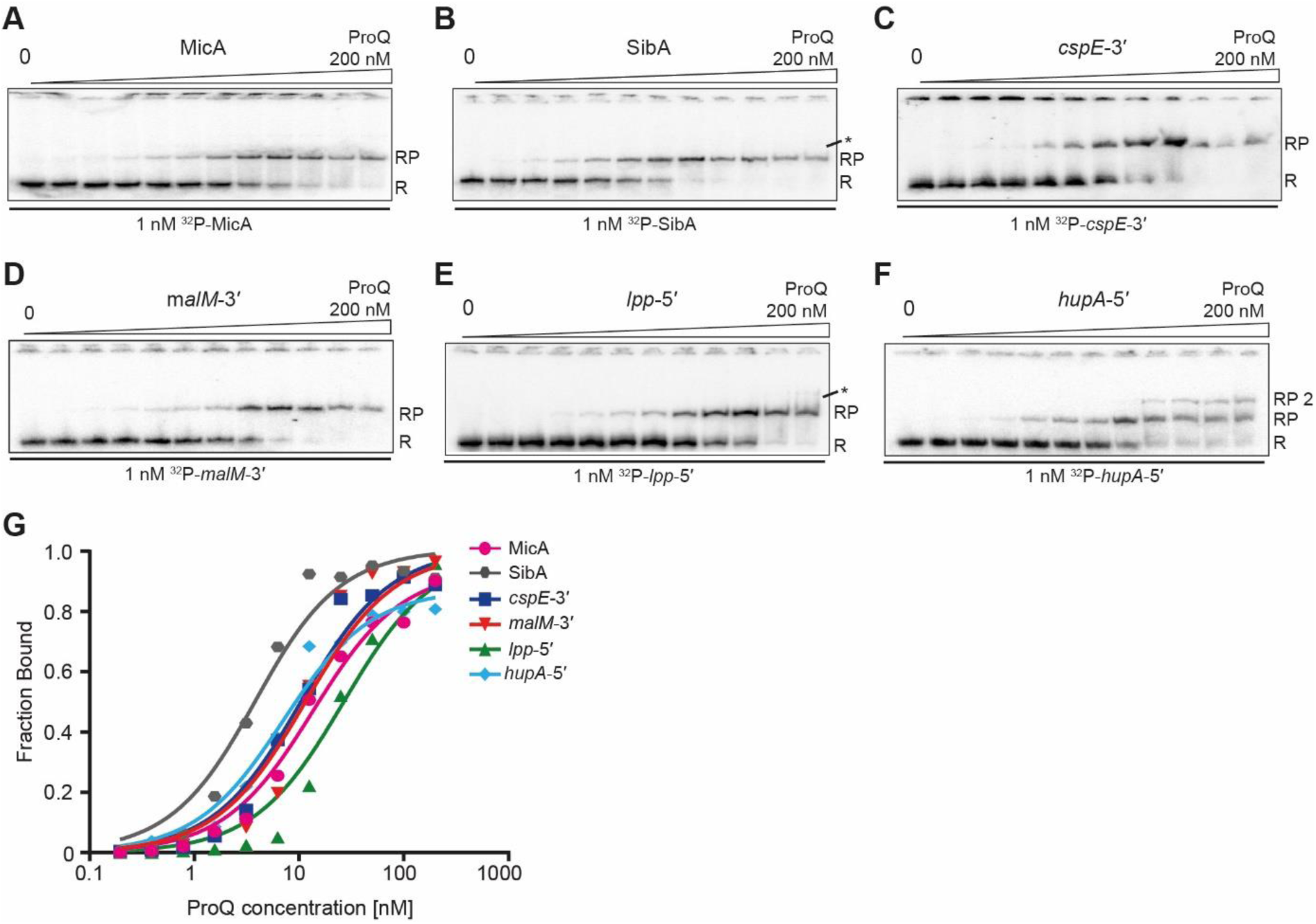
Equilibrium RNA binding to *E. coli* ProQ protein. The binding of ^32^P-labeled RNAs MicA (A), SibA (B), *cspE*-3′ (C), *malM*-3′ (D), *lpp*-5′ (E) and *hupA*-5′ (F) to full-length ProQ protein was monitored using a gelshift assay. Free ^32^P-RNA is marked as R, RNA-ProQ complex as RP and a higher order complex of *hupA*-5′ binding to ProQ as RP2. Asterisk (*) indicates additional RP complexes that could not be quantified due to low signal-to-noise ratio. (G) The fitting of the data from A-F using the quadratic equation provided *K*_d_ values of 13 nM for MicA, 3.3 nM for SibA, 10 nM for *cspE*-3′, 14 nM for *malM*-3′, 26 nM for *lpp*-5′ and 10 nM for *hupA*-5′. Fractions bound in both complexes of *hupA*-5′ with ProQ were summed together for the *hupA*-5′ *K*_d_ calculations. The average equilibrium dissociation constant (*K*_d_) values calculated from at least three independent experiments are shown in Table 1.

### The isolated FinO domain of ProQ preferentially binds RNAs that contain Rho-independent terminators

Given that previous studies showed contributions of ProQ’s FinO domain to RNA binding (19,29,30), the binding of this isolated domain to the six RNAs was also examined (Fig. 3, Table 1, Suppl. Fig. S1). The truncated protein construct (NTD) contained the N-terminal 130-aa of *E. coli* ProQ, which corresponds to the FinO domain, but was devoid of the linker and the C-terminal Tudor domain of ProQ (19,29). In the concentration range examined, all six RNAs formed single complexes with the NTD. However, when the binding affinity of the isolated NTD to the six RNAs was compared, the data showed striking differences (Fig. 3, Table 1). The affinities of MicA, SibA and *malM*-3’ to the isolated NTD were similar to their affinities to the ProQ protein (Fig. 3A, B, D, G, Table 1), and the affinity of *cspE*-3’ was 3-fold tighter than to the ProQ protein (Fig. 3C, G, Table 1). In contrast, the binding of *lpp*-5’ and *hupA*-5’ did not achieve saturation up to 1 µM concentration of the NTD, and hence their affinities to the NTD were at least 50-fold and 100-fold, respectively, weaker than to the full-length ProQ (Fig. 3E, F, G, Table 1). The fact that *lpp*-5’ and *hupA*-5’ mRNA fragments were the only RNAs from this set that did not possess Rho-independent transcription terminators (Fig. 1) suggested that 3’-terminal hairpin structures could be required for tight RNA binding by the isolated FinO domain of ProQ.

**Figure 3.**
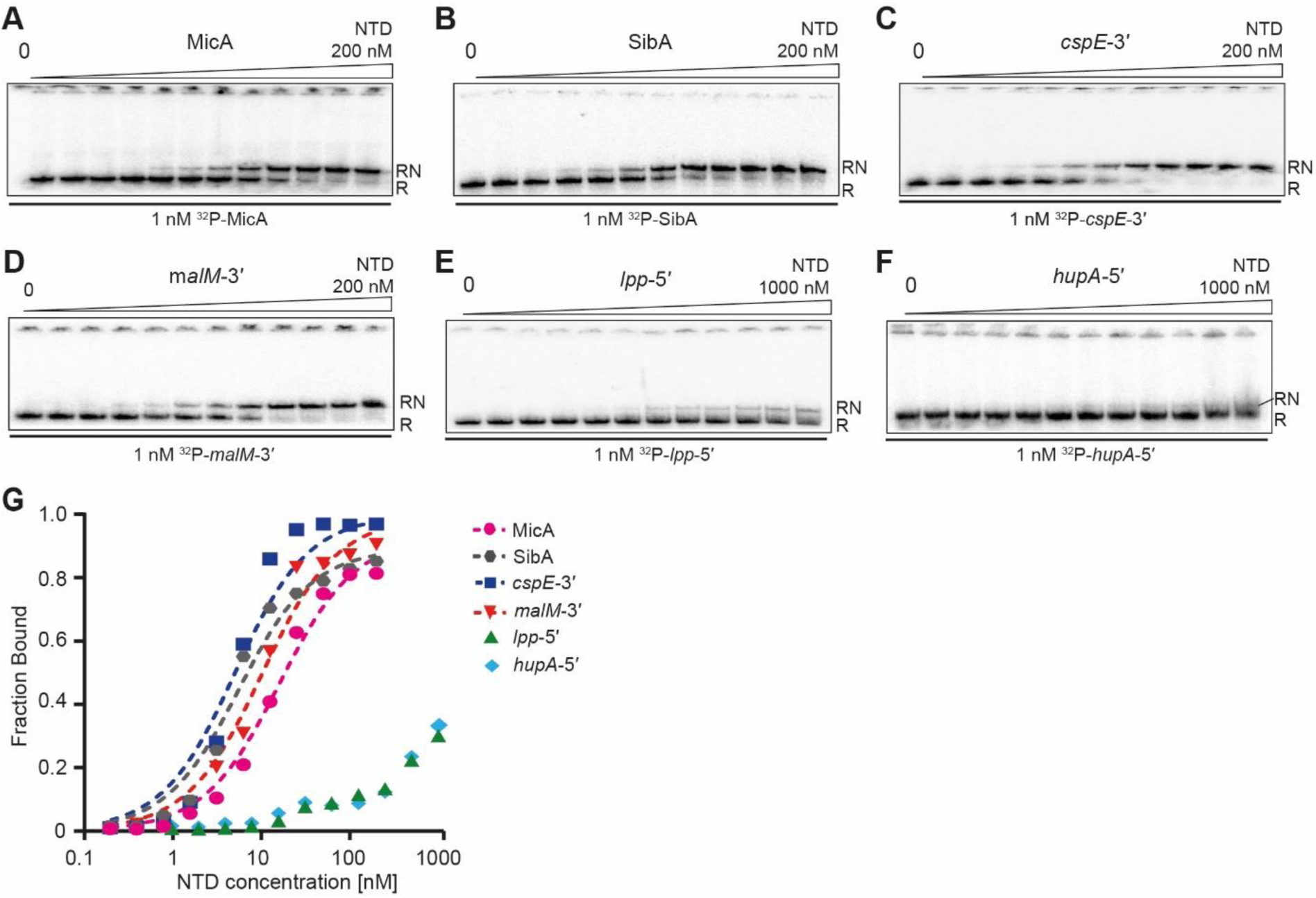
Equilibrium RNA binding to the isolated FinO domain of ProQ (NTD). The binding of 1 nM ^32^P-labeled RNAs MicA (A), SibA (B), *cspE*-3′ (C), *malM*-3′ (D), *lpp*-5′ (E) and *hupA*-5′ (F) to the NTD was monitored using a gelshift assay. Free ^32^P-RNA is marked as R, and RNA-NTD complex as RN. (G) The fitting of the data from A-F using the quadratic equation provided *K*_d_ values of 15 nM for MicA, 5 nM for SibA, 4.5 nM for *cspE*-3′, and 9.7 nM for *malM*-3′, while the affinities of *lpp*-5′ and *hupA*-5′ binding to the NTD were estimated as higher than 1 µM. The average *K*_d_ values obtained from at least three independent experiments are shown in Table 1.

### Binding by the ProQ NTD can be restored by the addition of a Rho-independent terminator structure

To test if the presence of terminator-like motifs on the 3’ ends of *lpp*-5’ and *hupA*-5’ would improve their binding to ProQ’s FinO domain, several point mutations were introduced into their 3’ terminal sequences to create structures similar to Rho-independent terminators (*lpp*-5’-mut and *hupA*-5’-mut) (Fig. 4A, B). Since both RNAs were predicted to have unstable secondary structures on the 3’ end, we designed a minimal set of mutations that would stabilize the double-stranded regions (mainly by changing G-U pairs into G-C pairs) and append short oligoU tails on the 3’ termini. The data showed that the presence of stable terminator-like structures in the *lpp*-5’-mut and *hupA*-5’-mut constructs strongly increased their affinity to the NTD. As a result, both mutants bound with similar affinities to the NTD and to ProQ (Fig. 4C, Table 1, Suppl. Fig. S2). Overall, these data showed that adding a hairpin motif followed by a short single-stranded tail to the 3’ end of an RNA could confer on it the ability to be bound by the isolated FinO domain of ProQ.

**Figure 4.**
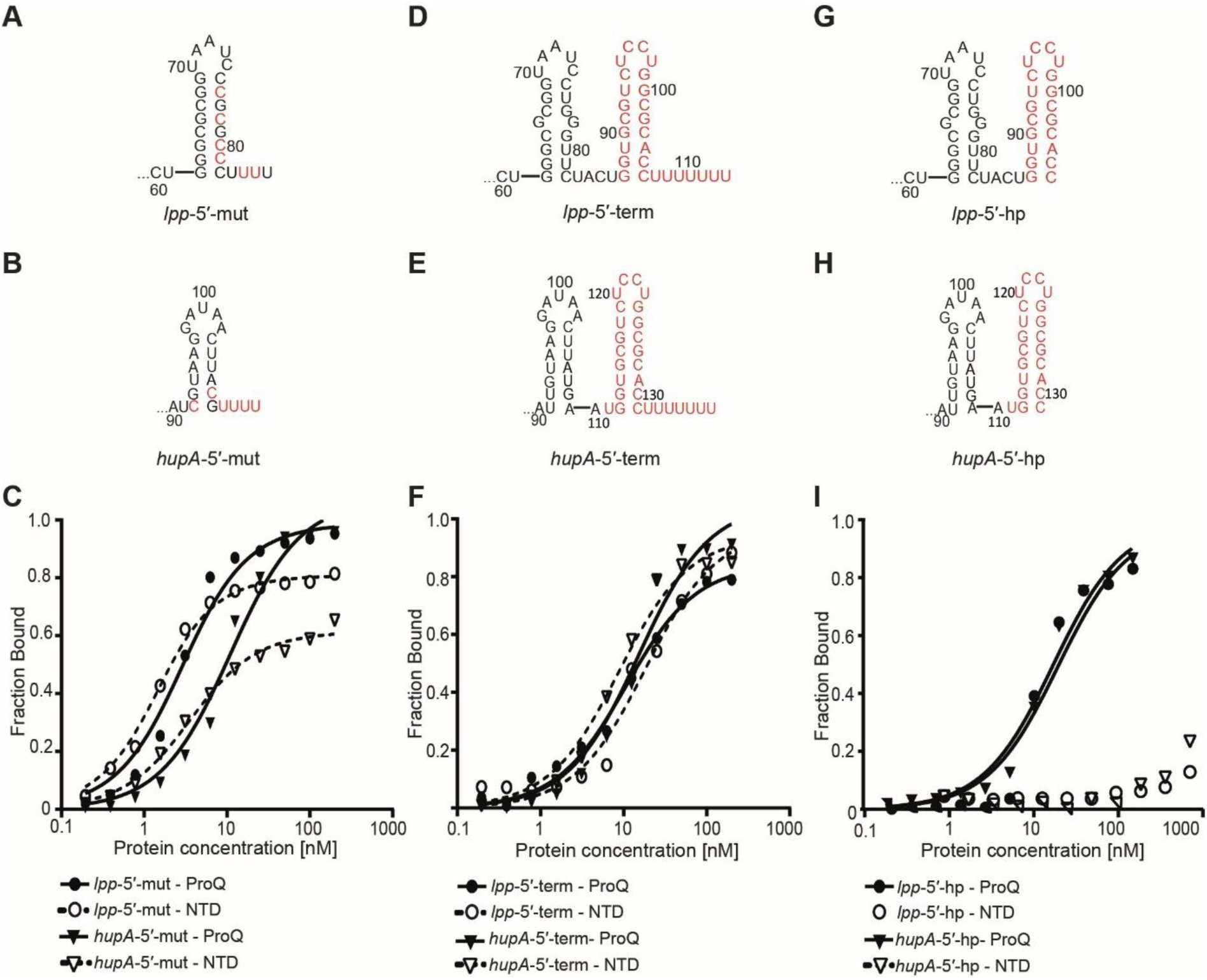
The Rho-independent terminators and similar structure motifs located on RNA 3′ ends are recognized by the FinO domain of ProQ. On top are presented the sequences of *lpp*-5′ mut (A), *hupA*-5′-mut (B), *lpp-*5′-term (D), *hup-*5′-term (E), *lpp*-5′-hp (G) and *hupA-*5′-hp (H) RNAs with point mutations and transplanted regions shown in red. The respective binding data with ProQ and the NTD are shown on graphs below each pair of mutants, in panels (C), (F), and (I), respectively. The fitting of the data using the quadratic equation provided *K*_d_ values of (C) 3.1 nM for *lpp*-5′-mut binding to ProQ and 7.6 nM to NTD, and 10 nM for *hupA*-5′-mut binding to ProQ and 18 nM to NTD; (F) 3 nM for *lpp*-5′-term binding to ProQ and 4.7 nM to NTD, and 15 nM for *hupA*-5′-term binding to ProQ and 8.2 nM to NTD; (I) 22 nM for *lpp*-5′-hp and 6.2 nM for *hupA*-5′-hp binding to ProQ. The binding of *lpp*-5′-hp and *hupA*-5′-hp to the NTD was essentially not detected up to 1 µM NTD concentration. Fractions bound in both complexes of *hupA*-5′-mut, *lpp*-5′-hp, and *hupA*-5′-hp with ProQ were summed together for the *K*_d_ calculations. Gels corresponding to the data in the plots are shown in Suppl. Fig. S2. Average *K*_d_ values are shown in Table 1.

To test if a natural Rho-independent terminator could also restore the binding to the NTD, the intrinsic terminator structure from *malM* 3’-UTR was transplanted to the 3’ ends of *lpp*-5’ and *hupA*-5’ to create *lpp*-5’-term and *hupA*-5’-term constructs (Fig. 4D, E). Binding data showed that *lpp*-5’-term and *hupA*-5’-term bound the NTD similarly to ProQ, as did *lpp*-5’-mut and *hupA*-5’-mut constructs (Fig. 4C, F, Table 1, Suppl. Fig. S2). Additionally, 3’-terminal oligoU sequences were removed from the transplanted terminators to leave only double-stranded motifs on the 3’ ends, thus creating *lpp*-5’-hp and *hupA*-5’-hp constructs (Fig. 4G, H). The binding of *lpp*-5’-hp and *hupA*-5’-hp to the NTD was essentially undetectable up to 1 µM concentration of the NTD (Fig. 4I, Table 1, Suppl. Fig. S2), which suggests that the 3’ oligoU sequence contributes to the RNA binding of the NTD. In summary, these data showed that adding a natural intrinsic terminator hairpin with a 3’ terminal oligoU sequence to the 3’ ends of *lpp*-5’ and *hupA*-5’ RNAs enabled their binding by the ProQ’s FinO domain.

To test if the location of the Rho-independent terminator structures at the very 3’ ends is required for their recognition by the ProQ NTD, the sequences of *cspE*-3’ and *malM*-3’ RNAs, which bind tightly to the NTD (Table 1), were extended beyond their natural 3’ ends (Fig. 5, Table 1, Suppl. Fig. S3). Each of these molecules was extended by either a shorter or longer length, resulting in four constructs named the *cspE*-3’-ext+5, *cspE*-3’-ext+22, *malM*-3’-ext+5, and *malM*-3’-ext+17 (Fig. 5). All four constructs bound strongly to ProQ with < 3-fold differences in binding affinities. Strikingly, however, these constructs differed markedly in binding to the isolated NTD (Fig. 5). The *cspE*-3’-ext+5 construct, which was extended by 5 nucleotides, bound the NTD only 3-fold weaker than *cspE*-3’ molecule. On the other hand, extension of *malM*-3’ by 5 nt showed a strong inhibitory effect, because the binding was not detected up to 1 µM concentration of the NTD. Similarly, the binding of both longer constructs, *cspE*-3’-ext+22 and *malM*-3’-ext+17, to the NTD was essentially undetectable up to 1 µM concentration of the NTD (Fig. 5, Table 1, Suppl. Fig. S3). In summary, the shorter extension had a detrimental effect only on the NTD binding of *malM*-3’ RNA, while the longer extensions were detrimental for the binding of both RNAs. One possibility is that extended sequences could form new secondary structures, potentially restricting the flexibility of the 3’ oligoU tails or making them less available for binding. This suggests that it is not simply the presence of a stretch of uridines, and/or their single-stranded nature, but also their presence at the 3’ terminus of the RNA that is important for binding to the NTD.

**Figure 5.**
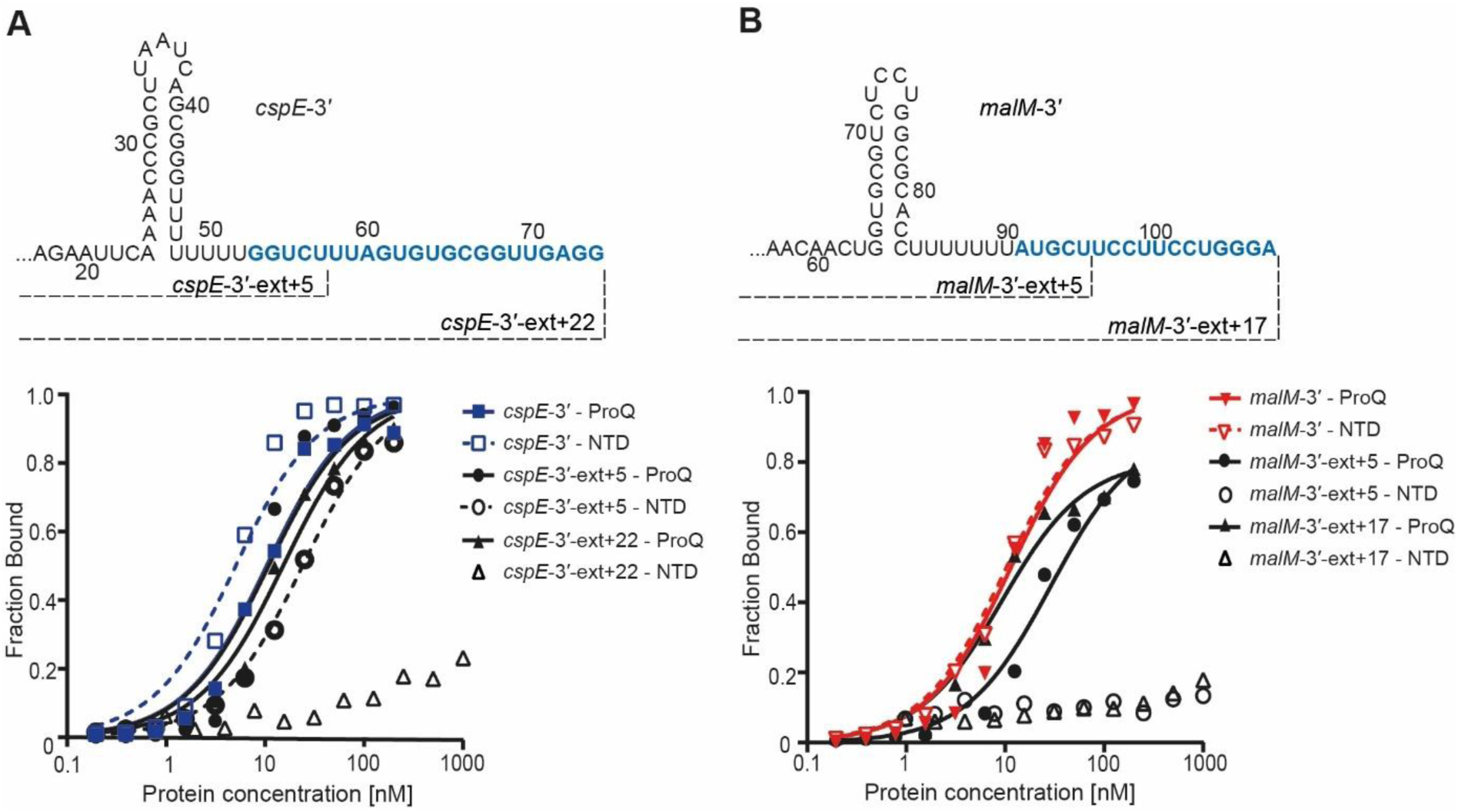
The extension of RNA sequence beyond the intrinsic terminator structure is detrimental for RNA binding by the FinO domain of ProQ. The sequences of *cspE*-3’ and *malM*-3′ were extended beyond the transcription terminators to obtain: (A) *cspE*-3′-ext+5 and *cspE*-3′-ext+22, and (B) *malM*-3′-ext+5 and *malM*-3′-ext+17 RNAs. The 3’ added sequences are shown in dark blue. The fitting of data using the quadratic equation provided *K*_d_ values of 13 nM for *cspE*-3′-ext+5 binding to ProQ and 24 nM for binding to NTD, 16 nM for *cspE*-3′-ext+22 binding to ProQ, 29 nM for *malM*-3′-ext+5 binding to ProQ and 10 nM for *malM*-3′-ext+17 binding to ProQ, while *cspE*-3′-ext+22, *malM*-3′-ext+5 and *malM*-3′-ext+17 binding to NTD was essentially undetectable up to 1 µM concentration of the protein. Fractions bound in both complexes of *malM*-3′-ext+17 with full-length ProQ were summed together for *K*_d_ calculations. The data in the plots for *cspE*-3′ and *malM*-3′ binding to ProQ or NTD are the same as in Figures 2 and 3, respectively. Gels corresponding to the data in the plots are shown in Suppl. Fig. S3. Average *K*_d_ values are shown in Table 1.

### The double-stranded region in the lower part of the terminator hairpin is essential for tight RNA binding by the FinO domain

Several recent studies showed that ProQ binding sites in *E. coli* and *S. enterica* transcriptomes were enriched in double-stranded RNA structural elements (11-13). To better understand the contribution of the *malM*-3’ terminator hairpin to the FinO domain binding, three constructs with mutations that interrupt the continuous double-stranded region in the middle part of this hairpin were designed (Fig. 6A). The results showed that a triple-adenosine bulge and double A-A and U-U mismatches, each inserted in the middle of the stem, did not have detrimental effects on the binding to the NTD (Fig. 6A, C, Table 2, Suppl. Fig. S4). This could suggest that in each case the disturbed region of the helical structure was located outside of the region of the terminator stem necessary for the binding to the NTD.

**Table 2.**
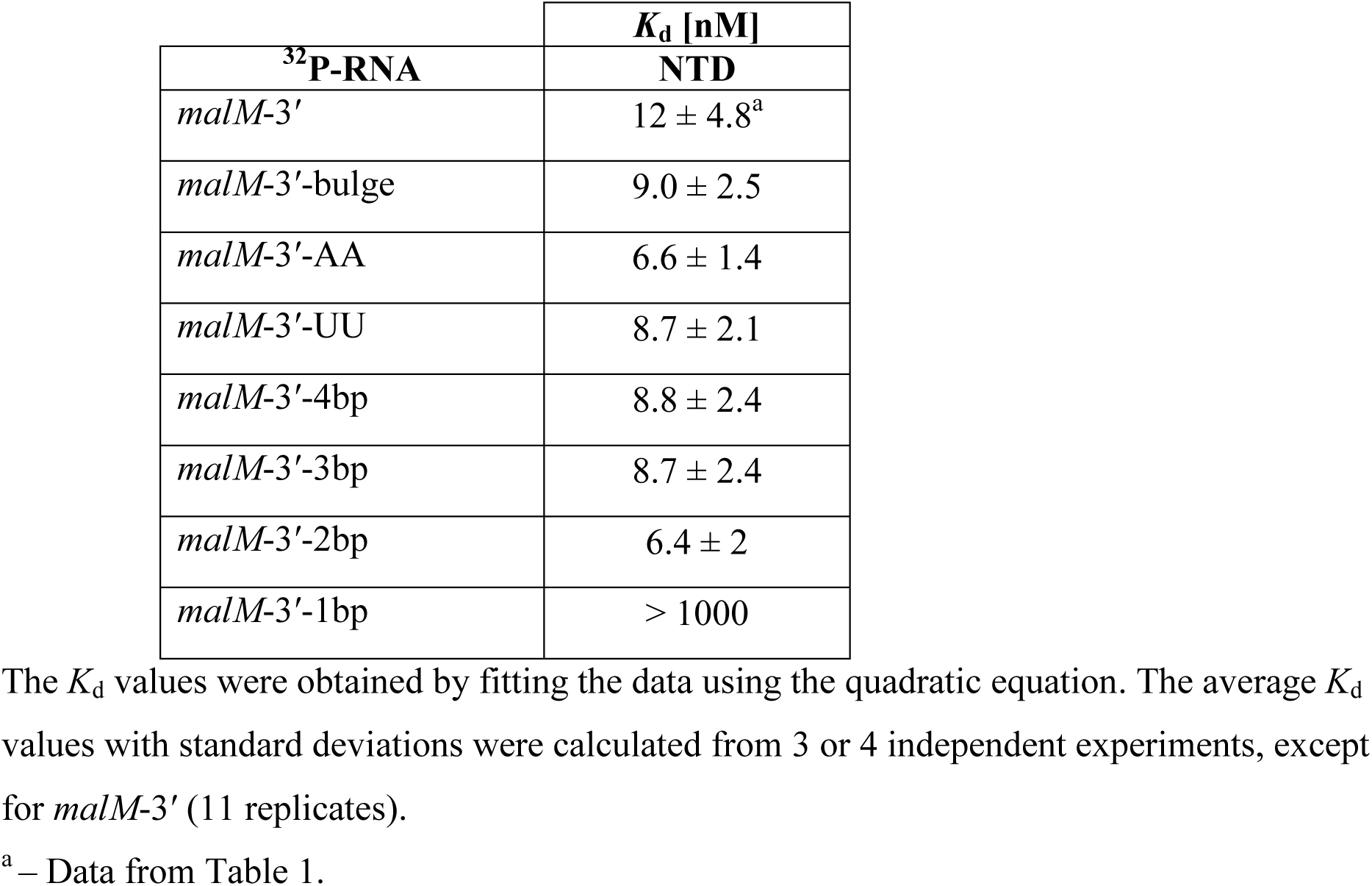
Short double-stranded region of the 3′-terminal hairpin of *malM*-3′ RNA is sufficient for binding by the ProQ NTD.

| <sup>32</sup>P-RNA | *K<sub>d</sub>* [nM] |
| --- | --- |
|  | NTD |
| *malM*-3' | 12 ± 4.8<sup>a</sup> |
| *malM*-3'-bulge | 9.0 ± 2.5 |
| *malM*-3'-AA | 6.6 ± 1.4 |
| *malM*-3'-UU | 8.7 ± 2.1 |
| *malM*-3'-4bp | 8.8 ± 2.4 |
| *malM*-3'-3bp | 8.7 ± 2.4 |
| *malM*-3'-2bp | 6.4 ± 2 |
| *malM*-3'-1bp | > 1000 |

The *K<sub>d</sub>* values were obtained by fitting the data using the quadratic equation. The average *K<sub>d</sub>* values with standard deviations were calculated from 3 or 4 independent experiments, except for *malM*-3' (11 replicates).
<sup>a</sup> – Data from Table 1.

**Figure 6.**
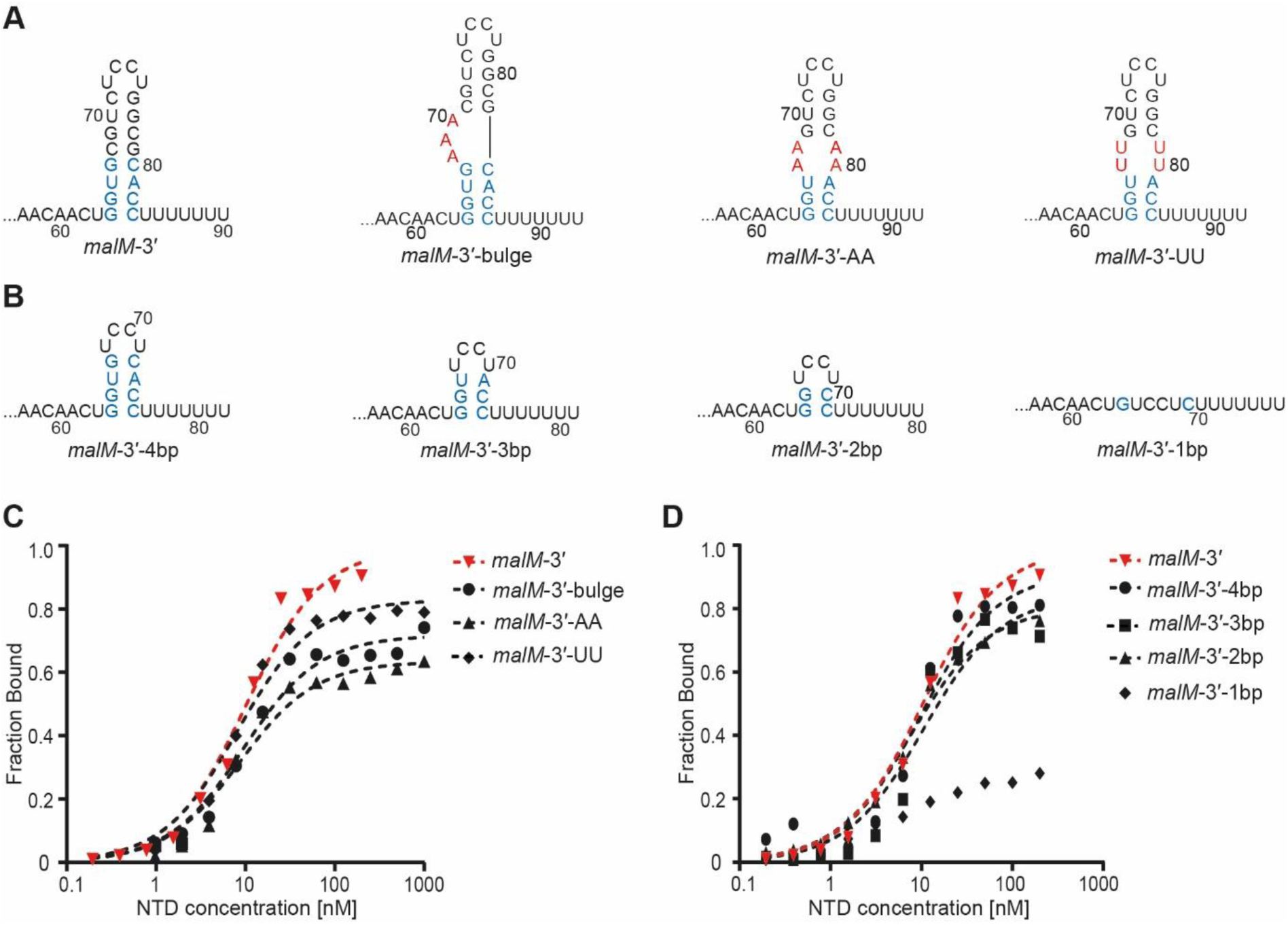
Short double-stranded stem of the terminator structure is sufficient for *malM*-3′ binding to the FinO domain. The nucleotides involved in the four base-pairs at the bottom of the terminator stem are shown in light blue font, and the mutations introduced into *malM*-3′-bulge, *malM*-3′-AA and *malM*-3′-UU RNAs are shown in red font. The mutants with shorter stems were constructed by gradual removal of the base-paired nucleotides from the top part of the terminator stem. The fitting of the binding data in the plot using the quadratic equation provided *K*_d_ values of 8.9 nM for *malM*-3′-bulge, 8.2 nM for *malM*-3′-AA, 7.6 nM for *malM*-3′-UU, 9.1 nM for *malM*-3′-4bp, 10 nM for *malM*-3′-3bp, 7.2 nM for *malM*-3′-2bp, while the binding of *malM*-3′-1bp construct did not reach saturation up to 1µM concentration of the NTD. Data for wt *malM*-3′ are the same as in Figure 3D. Gels corresponding to the data in the plots are shown in Suppl. Fig. S4. Average *K*_d_ values are shown in Table 2.

To test how the length of continuous base-pairing in the terminator stem could affect the binding, additional constructs with shorter stems were designed (Fig. 6B). These constructs were made in such a way that the base-paired nucleotides were gradually removed from the top part of the *malM*-3’ terminator stem, while the apical loop and base-paired nucleotides at the bottom of the stem remained. Binding data showed that those constructs, which were predicted to have 4, 3 or 2 base pairs of the stem remaining, bound the NTD with similar affinities as the wt *malM*-3’. However, the construct possessing only the nucleotides corresponding to the bottom base pair of the stem, did not bind to the NTD (Fig. 6B, D, Table 2, Suppl. Fig. S4). As the nucleotide sequence corresponding to the stem of this construct would likely be unfolded, this is consistent with a stably base-paired structure being necessary for the NTD binding. In summary, the effects of deleting predicted base pairs and inserting disruptions into the double-stranded region of the RNA terminator hairpin together suggest that only 2-3 continuous base pairs preceding the U-stretch supports strong binding to the isolated NTD domain.

### Shortening the 3’-terminal oligoU sequence to less than four uridine residues abolishes binding by the FinO domain of ProQ

It was recently observed that 3’ oligoU sequences were part of the sequence logo of ProQ binding sites detected by RIL-seq in *E. coli* (12), and that 3’ oligoU tails of RNA ligands of ProQ were shorter than those of RNA ligands of Hfq in *S. enterica* (11). To elucidate the role of the 3’ oligoU sequence in binding to the NTD of ProQ, versions of *malM*-3’ were designed with varying lengths of 3’ oligoU tails (Fig. 7). The natural terminator of *malM*-3’ contains seven 3’-terminal uridine residues (Fig. 1). Extending the 3’-oligoU sequence of *malM*-3’ by two uridines in the *malM*-3’-9U construct had only a small effect on the binding affinity to the NTD (Fig. 7, Table 3, Suppl. Fig. S5). On the other hand, a series of constructs with truncations of the 3’ oligoU sequence resulted in a range of effects on the NTD binding. The constructs with five or six uridines remaining bound the ProQ NTD similarly to wt *malM*-3’. When one additional uridine residue was removed, the resulting *malM*-3’-4U construct bound the NTD approximately 2-fold weaker than the wt *malM*-3’. Even more dramatic effects were observed when an additional uridine was removed: the binding of *malM*-3’-3U construct, which had only three uridine residues in the 3’ tail, did not reach saturation up to 1 µM concentration of the NTD, while the binding of the constructs with even shorter tails was essentially undetectable (Fig. 7, Table 3, Suppl. Fig. S5). Hence, at least four 3’-terminal uridine residues were necessary for the strong binding of *malM*-3’ to the isolated FinO domain of ProQ.

**Table 3.** The length of single-stranded sequence following the Rho-independent terminators affects the binding of *malM*-3′ RNA to the N-terminal domain of ProQ.

| | $K_d$ [nM] |
| --- | --- |
| <sup>32</sup>P-RNA | NTD |
| *malM*-3'-9U | $5.1 \pm 2.9$ |
| *malM*-3' (7U, wt) | $12 \pm 4.8^a$ |
| *malM*-3'-6U | $7.8 \pm 1.2$ |
| *malM*-3'-5U | $9.7 \pm 1.5$ |
| *malM*-3'-4U | $27 \pm 11$ |
| *malM*-3'-3U | $> 1000$ |
| *malM*-3'-2U | n. d. |
| *malM*-3'-1U | n. d. |

The $K_d$ values were obtained by fitting the data to the quadratic equation. The average $K_d$ values with standard deviations were calculated from 3 to 5 independent experiments, except for *malM*-3' (11 replicates).
<sup>a</sup> – Data from Table 1.
n. d. – the binding was essentially undetectable up to 1 μM concentration of the NTD.

**Figure 7.**
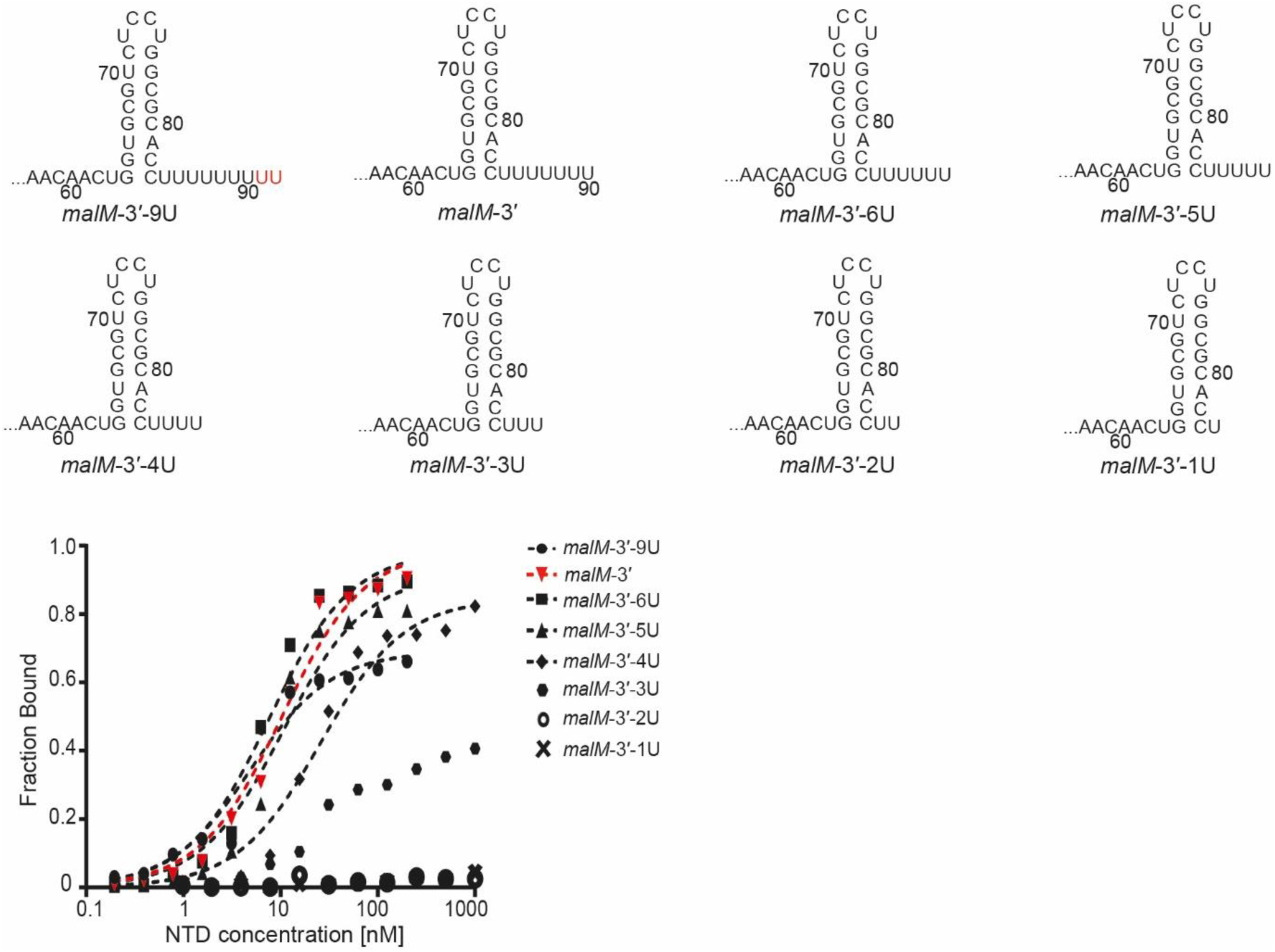
The binding of *malM*-3′ constructs with different lengths of 3’-terminal oligoU tails to the FinO domain. Nucleotides added on 3’ end of *malM*-3′-9U are marked with red font. The fitting of data shown on the graph using the quadratic equation provided *K*_d_ values of 4.4 nM for *malM*-3′-9U, 7.1 nM for *malM*-3′-6U, 10 nM for *malM*-3′-5U and 25 nM for *malM*-3′-4U, while the binding of *malM*-3′-3U did not reach saturation up to 1 µM concentration of the NTD. The binding of *malM*-3′-2U and *malM*-3′-1U was essentially undetectable up to 1 µM concentration of the NTD. Data for *malM*-3′ are the same as in Figure 3 D. Gels corresponding to the data in the plots are shown in Suppl. Fig. S5. Average *K*_d_ values are shown in Table 3.

To corroborate the observation that the length of the 3’ oligoU sequence affects the RNA binding by the ProQ NTD, three constructs of *cspE*-3’ with truncated 3’ oligoU stretches were compared to the wt RNA (Suppl. Fig. S6). The intrinsic terminator structure of *cspE*-3’ contains eight uridine residues in the 3’-oligoU sequence, with four of these residues predicted to be base-paired (Fig. 1). When two 3’-terminal uridine residues were removed, the resulting construct *cspE*-3’-6U bound the NTD with the same affinity as the wt *cspE*-3’. However, when two more uridine residues were removed, the *cspE*-3’-4U construct, in which the remaining part of the 3’-tail was fully base-paired, bound the NTD about 10-fold weaker than the wt *cspE*-3’. Additionally, when the entire oligoU sequence was removed in the *cspE*-3’-0U construct, binding was fully abolished (Suppl. Fig. S6). These data support the conclusion that the minimum length of four uridine residues of the 3’-oligoU sequence is necessary for RNA binding by the NTD. The fact that the binding of *cspE*-3’-4U was weaker than the binding of *malM*-3’-4U could suggest that because the 3’-terminal uridines of *cspE*-3’-4U are base-paired with adenosine residues on the 5’ side of the terminator, they are less available for binding by the ProQ NTD.

### RNAs bound by ProQ are enriched for A-rich sequences on the 5’ side of the intrinsic terminator

Binding of the ProQ NTD to intrinsic terminator structures (Figs. 3-5) is reminiscent of the Hfq protein, which also binds the terminator hairpins with 3’ oligoU tails (39). This poses an interesting question of how these proteins discriminate between their RNA ligands, because the ProQ and Hfq-specific RNAs constitute mostly separate RNA pools in bacterial cells (11-13). One hint comes from the recent observation that RNA ligands of ProQ identified by CLIP-seq had significantly shorter single-stranded lengths of 3’ oligoU tails, which was caused by their partial involvement in base-pairing with nucleotide residues on the 5’ side of the hairpin stem (11). Another hint comes from the fact that an U-rich sequence 5’ adjacent to the Rho-independent terminator stem is a part of the Hfq-binding module in some of its RNA ligands, predominantly *trans*-encoded sRNAs (39). These observations led us to hypothesize that the sequences on the 5’ side of the terminator hairpin could contribute to discrimination between RNA molecules by the ProQ and Hfq proteins.

When we examined the sequences of ProQ-specific RNAs in previously reported datasets (11,12), we noticed that they contained an A-rich sequence on the 5’ side of the terminator (Fig. 8, Suppl. Fig. S7, Suppl. Table S3). We began by computing sequence logos for a set of the top 50 terminator-containing RNAs, either mRNA 3’ UTRs or sRNAs, which were most enriched in the RIL-seq data of either ProQ- or Hfq-binding RNAs in *E. coli* (12). We analyzed the nucleotide frequency in a 10-nt long sequence on the 5’ side of the terminator of these RNAs. In this region close to the terminator stem, adenosines were overrepresented in the ProQ-specific RNAs (Fig. 8A, Suppl. Table S3). This A-rich motif was also present in ProQ-specific RNAs identified in *E. coli* using CLIP-seq (Suppl. Fig. S7 A, Suppl. Table S3) (11). In contrast, in Hfq-specific RNAs, uridines were most common in the corresponding sequence (Fig. 8B, Suppl. Table S3). In further support, adenosines were significantly overrepresented when ProQ-specific RNAs were directly compared to Hfq-specific RNAs (Fig. 8C) (12,48). In the datasets obtained in *S. enterica* using CLIP-seq, adenosines were also overrepresented in this region of ProQ-specific RNAs in comparison to Hfq-specific RNAs (Suppl. Fig. S7 B-D, Suppl. Table S3) (11). In summary, the results of this analysis showed that an A-rich motif on the 5’ side of the terminator is a sequence feature present in ProQ-specific RNAs but not Hfq-specific RNAs.

**Fig. 8.**
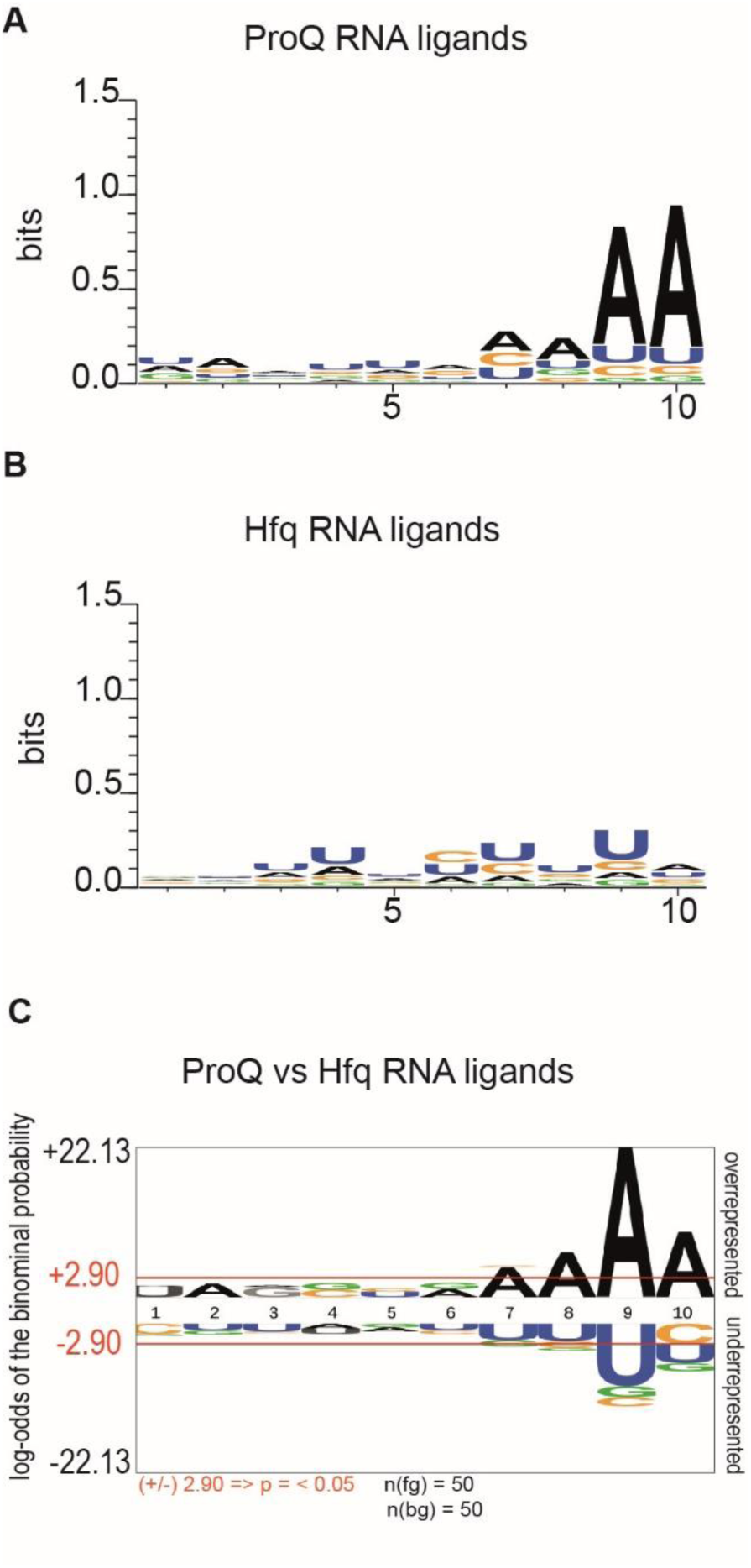
The sequence on the 5’ side of the intrinsic terminator hairpin in top RNA ligands of ProQ is enriched in adenosine residues. Nucleotide frequencies in the 10-nt long sequence 5’ adjacent to the terminator hairpin are shown for top 50 RNA ligands of ProQ and Hfq containing Rho-independent terminators, in the data obtained previously by RIL-seq in *E. coli* (12). (A) Nucleotide frequencies obtained by *WebLogo* software for the top 50 *E. coli* ProQ ligand RNAs containing Rho-independent terminators in RIL-seq data, p < 2.2 × 10^−16^; (B) Nucleotide frequencies obtained by *WebLogo* for the top 50 *E. coli* Hfq ligand RNAs containing Rho-independent terminators in RIL-seq data, p = 6.5 × 10^−12^; (C) The statistically significant nucleotide frequencies at each nucleotide position for ProQ RNA ligands as compared to Hfq RNA ligands. Sequences were analyzed by *pLogo* software for top 50 *E. coli* ProQ RNA ligands as the foreground, and top 50 Hfq target RNAs as the background. Statistically significant p-values: A at position 7, p = 0.002; A at position 8, p = 1.08 × 10^−5^; A at position 9, p = 2.99 × 10^−21^; A at position 10, p = 9.78 × 10^−9^. The number of foreground sequences was marked on the figure as n(fg), and the number of background sequences as n(bg).

### The A-rich motif discourages the RNA binding to the Hfq protein

To determine the role of the A-rich motif in discriminating between ProQ and Hfq binding, this sequence was mutated to uridines in several RNAs, and their binding to the ProQ NTD and to the Hfq protein were compared (Figs. 9-111, Suppl. Figs. S8-S10). For this analysis we selected such molecules that were at least 60-fold enriched in RIL-seq data for *E. coli* (12), that had well defined A-rich motifs, and for which the A-to-U substitutions were not predicted to have large-scale effects on secondary structure, as predicted by *RNAStructure* software. In addition to *malM*-3’ and *cspE*-3’, RNAs selected for the analysis were *cspA-*3’, *hupB-*3’, *infA*-3’, *ihfA*-3’, and *gapA*-3’ (Fig. 9, Suppl. Fig. S9A, B). Because recent bacterial three-hybrid experiments used a longer sequence of *cspE*-3’ (30), we also analyzed a longer variant of this molecule, named *cspE*81-3’. We substituted a continuous stretch of nucleotides, which in two cases also included single guanosines predicted to base pair with uridines of the 3’ oligoU tail, for all of the RNAs except *malM*-3’ where the substitution was interrupted by one nucleotide (Fig. 9).

**Figure 9.**
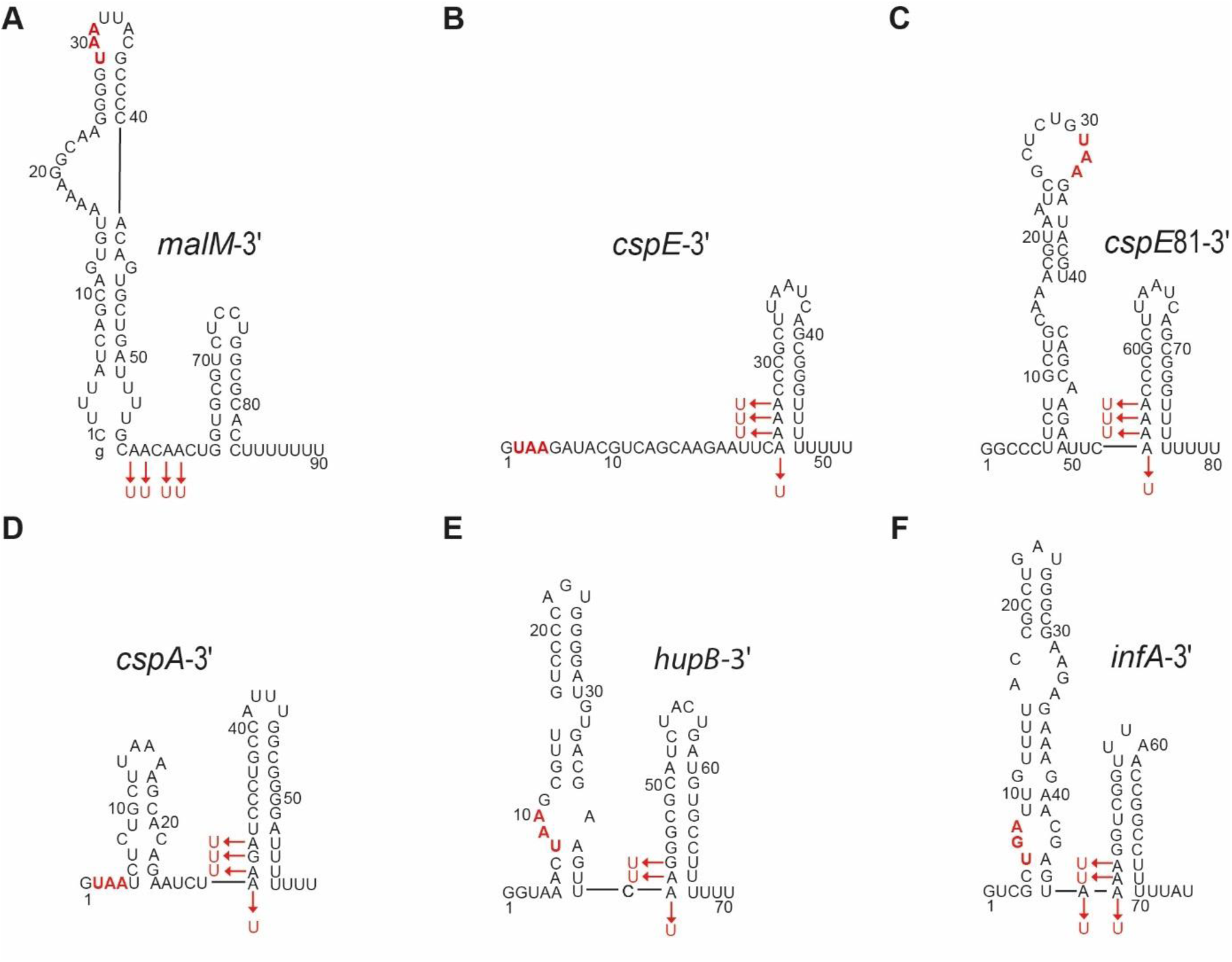
The mutations introduced into RNA molecules to test the role of A-rich motifs on the 5’ side of the terminator hairpins. The secondary structures of *malM*-3’, *cspE*-3’, *cspE*81-3’, *cspA*-3’, *hupB*-3’, and *infA*-3’ RNAs with mutations introduced into the A-rich regions marked with red font. The stop codons are marked in bold red font. The lower case g denotes guanosine residue added on 5′ end of *malM*-3′ and *malM*-3′-AtoU sequences to enable T7 RNA polymerase transcription.

The substitutions of the A-rich motif to uridines had either small effects or were detrimental for the binding of mutant RNAs to the ProQ NTD (Fig. 10, Table 4, Suppl. Figs. S8, S9). The mutations in *malM*-3’, *cspA-*3’, *infA*-3’, *hupB-*3’, and *gapA*-3’ had 2-fold or smaller effects on the binding affinity to the NTD. On the other hand, mutations in both variants of *cspE*-3’ and in *ihfA*-3’ had more than 10-fold detrimental effects on binding to the NTD (Table 4, Suppl. Fig. S9).

**Table 4.** Equilibrium binding of wt RNAs and their variants with mutations in the A-rich motifs to the ProQ NTD and the Hfq protein.

| <sup>32</sup>P-RNA | $K_d$ [nM] | |
| --- | --- | --- |
|  | NTD<sup>a</sup> | Hfq<sup>b</sup> |
| *malM*-3' | 12 ± 4.8<sup>c</sup> | 22 ± 7 |
| *malM*-3'-AtoU | 5.0 ± 1.8 | 2.2 ± 0.8 |
| *cspE*-3' | 4.5 ± 1.5<sup>c</sup> | 5.9 ± 2.5 |
| *cspE*-3'-AtoU | 100 ± 51 | 1.7 ± 0.88 |
| *cspE*81-3' | 7.6 ± 4.5 | 47 ± 11 |
| *cspE*81-3'-AtoU | 17 ± 4.6 | 2.6 ± 1.2 |
| *cspA*-3' | 33 ± 10<sup>d</sup> | 49 ± 15<sup>d</sup> |
| *cspA*-3'-RtoU | 42 ± 18<sup>d</sup> | 1.3 ± 0.66 |
| *hupB*-3' | 18 ± 1.7 | 30 ± 16<sup>d</sup> |
| *hupB*-3'-RtoU | 29 ± 7.2 | 1.9 ± 0.75 |
| *infA*-3' | 19 ± 10 | 28 ± 12<sup>d</sup> |
| *infA*-3'-AtoU | 32 ± 13 | 1.2 ± 0.55 |

The $K_d$ values were obtained by fitting the data to the quadratic equation. The average $K_d$ values with standard deviations were calculated from 3 to 5 independent experiments, except for *malM*-3' binding to the NTD (11 replicates) and Hfq (6 replicates), and *cspE*-3' binding to the NTD (7 replicates) and Hfq (7 replicates).
<sup>a</sup> – The RNA binding to the NTD was measured using gelshift assay
<sup>b</sup> – The RNA binding to the Hfq protein was measured using double-filter retention assay
<sup>c</sup> – Data from Table 1.
<sup>d</sup> – The endpoint value of 100% was assumed in the data fitting.

**Figure 10.**
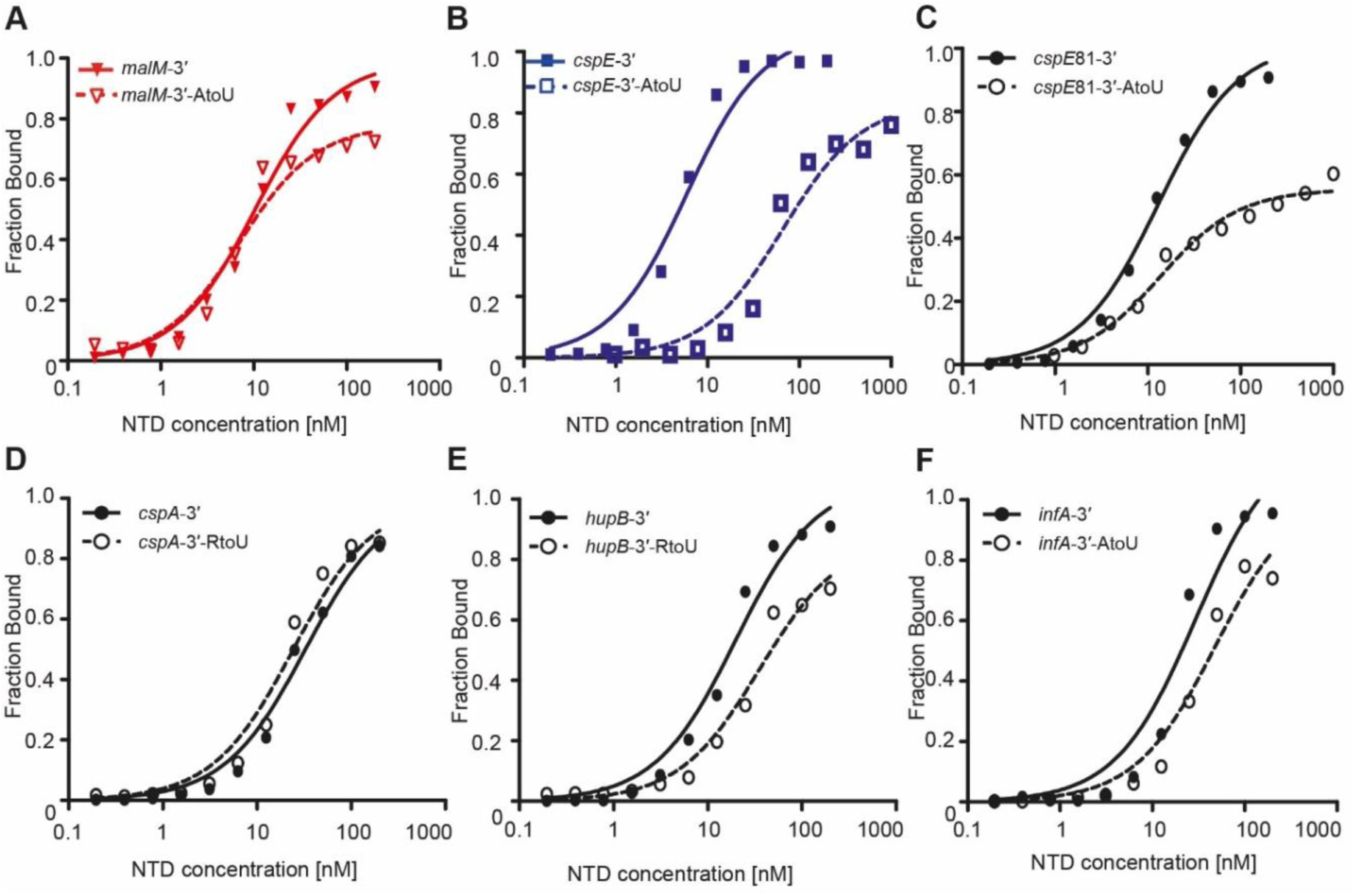
The binding of wt RNAs and their variants with mutations in the A-rich motifs to the FinO domain. The binding of ^32^P-labeled RNAs to the NTD was monitored using a gelshift assay. The fitting of the data using the quadratic equation provided *K*_d_ values of (A) 9.7 nM for *malM*-3′, 6.5 nM *malM*-3′-AtoU; (B) 4.5 nM for *cspE*-3′, 65 nM for *cspE*-3′-AtoU; (C) 12 nM for *cspE*81-3′, 13 nM for *cspE*81-3′-AtoU; (D) 32 nM for *cspA*-3′, 24 nM for *cspA*-3′-RtoU; (E) 19 nM for *hupB*-3′, 35 nM for *hupB*-3′-RtoU; (F) 27 nM for *infA*-3′, 47 nM for *infA*-3′-AtoU. Data for *cspE-*3′ and *malM*-3′ are the same as in Figure 3 C and D, respectively. Gels corresponding to the data in the plots are shown in Suppl. Fig. S8. The average equilibrium dissociation constant (*K*_d_) values calculated from at least three independent experiments are shown in Table 4.

In contrast to the neutral-to-negative effects on ProQ binding, substitutions of the A-rich motif to uridines strongly increased the affinities of these RNAs to the Hfq protein (Fig. 11, Table 4, Suppl. Figs. S9, S10). The binding of *E. coli* Hfq protein to the studied RNAs was measured using a double-filter retention assay, which was previously used to compare the binding of different sRNAs to the Hfq protein (40,44). The advantage of this assay is that the separation of the RNA-Hfq complex from free RNAs occurs rapidly, and it allows parallel comparison of up to 8 different RNAs (44). For all of the mutants, except *cspE*-3’-AtoU, the binding to Hfq was at least 10-fold tighter than to their wt counterparts. The strongest effects were observed for *cspA*-3’-RtoU, *ihfA*-3’-AtoU, and *gapA*-3’-AtoU, and smaller effects were seen for *malM*-3’-AtoU, *hupB-*3’-RtoU, and *infA*-3’-AtoU (Fig. 11, Table 4, Suppl. Fig. S9).

**Figure 11.**
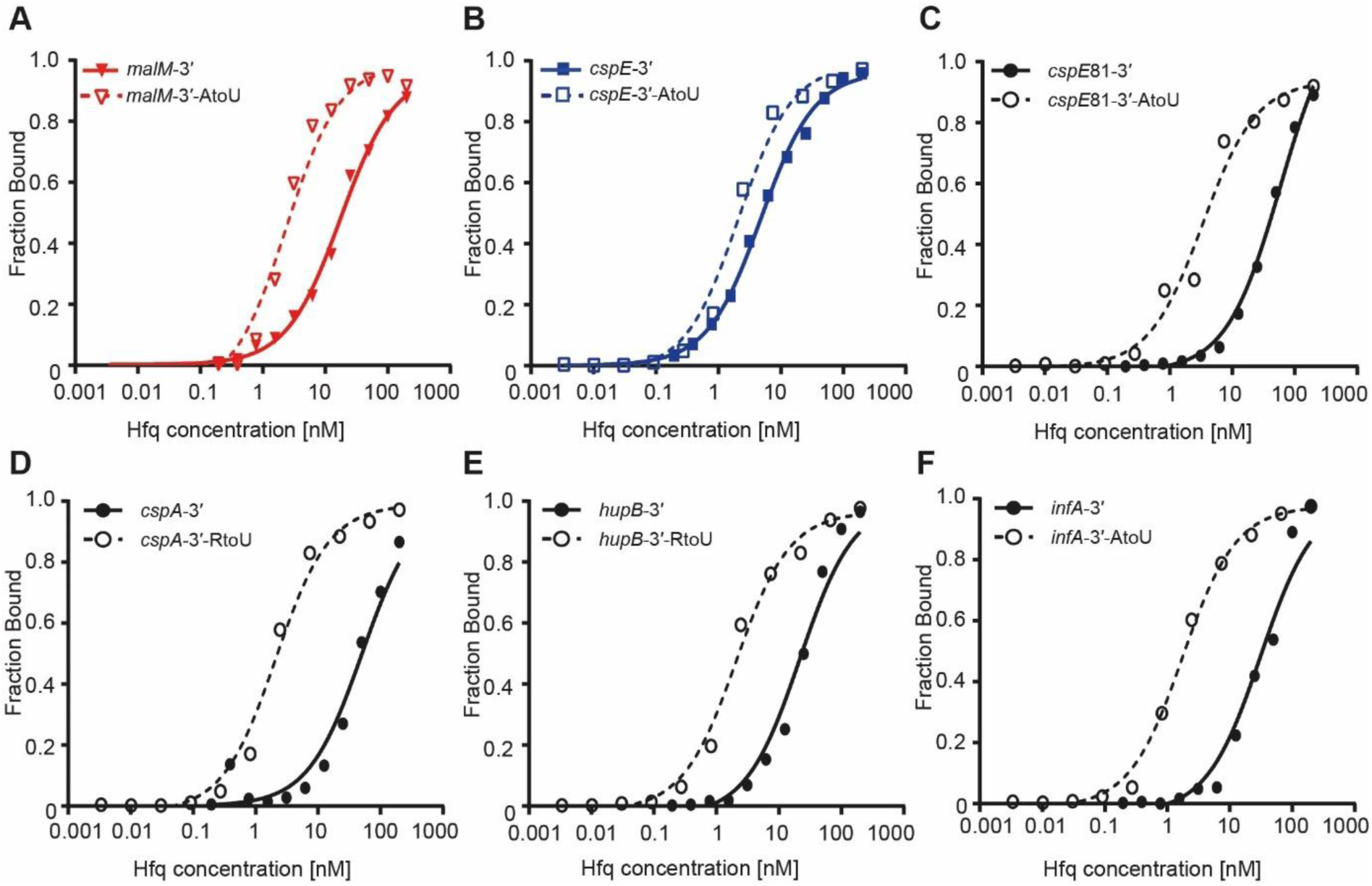
The binding of wt RNAs and their variants with mutations in the A-rich motifs to the *E. coli* Hfq protein. The binding of ^32^P-labeled RNAs to the Hfq protein was monitored using a double-filter retention assay. The fitting of the data using the quadratic equation provided *K*_d_ values of (A) 17 nM for *malM*-3’, 2.3 nM *malM*-3’-AtoU; (B) 4.7 nM for *cspE*-3′, 2.1 nM for *cspE*-3′-AtoU; (C) 40 nM for *cspE*81-3′, 2.8 nM for *cspE*81-3′-AtoU; (D) 51 nM for *cspA*-3′, 2.1 nM for *cspA*-3′-RtoU; (E) 22 nM for *hupB*-3′, 2 nM for *hupB*-3′-RtoU; (F) 31 nM for *infA*-3′, 1.7 nM for *infA*-3′-AtoU (F). Raw filter data are shown in Suppl. Fig. S10. The average equilibrium dissociation constant (*K*_d_) values calculated from at least three independent experiments are shown in Table 1.

While the binding of the *cspE*-3’-AtoU mutant to Hfq was not affected by the substitutions, the longer construct *cspE*81-3’-AtoU bound Hfq 18-fold tighter than its wt version (Fig. 11, Table 4, Suppl. Fig. S10). The wt *cspE*-3’ molecule has a long single-stranded sequence in its 5’ region, while the corresponding sequence is involved in secondary structure in the *cspE*81-3’ molecule (Fig. 9 B, C). Because this sequence could serve as an additional Hfq binding site in wt *cspE*-3’, it is likely that the effect of the A-to-U substitutions in *cspE*-3’ is masked by the Hfq binding to this additional sequence. Overall, these data showed that the substitution of adenosines to uridines on the 5’ side of the terminator hairpin had different effects on Hfq and ProQ NTD binding – increasing the affinity of the Hfq protein to these RNAs while at the same time not altering or decreasing the affinity of the ProQ NTD.

### *In vivo* RNA-binding determinants for ProQ and Hfq

The above results indicate that, in a purified *in vitro* system, adenosines immediately upstream of intrinsic terminators serve as negative determinants for Hfq binding (Figs. 9, 11, Table 4). In contrast, these adenosines appear to be neutral or detrimental factors for the ProQ NTD binding (Fig. 10). An important question is whether these adenosines play a similar role in controlling Hfq- and ProQ-interactions *in vivo*. To investigate this, we utilized a bacterial three-hybrid (B3H) system for RNA-protein interactions that has been demonstrated to detect both Hfq- and ProQ-RNA interactions *in vivo* (30,51). In this assay, expression of three hybrid components make reporter-gene (*lacZ)* transcription from a test promoter dependent on a productive interaction between an RNA ‘bait’ and a protein ‘prey’ (Fig. 12A). Specifically, a “Bait” RNA is expressed as an MS2^hp^-hybrid RNA, and is tethered to DNA upstream of the transcription start site by interaction between a CI-MS2^CP^ “Adapter” protein and the λ operator O_L_2 sequence. Prey protein is expressed as a fusion with the N-terminal domain of the alpha subunit (α) of RNA polymerase (RNAP).

**Figure 12.**
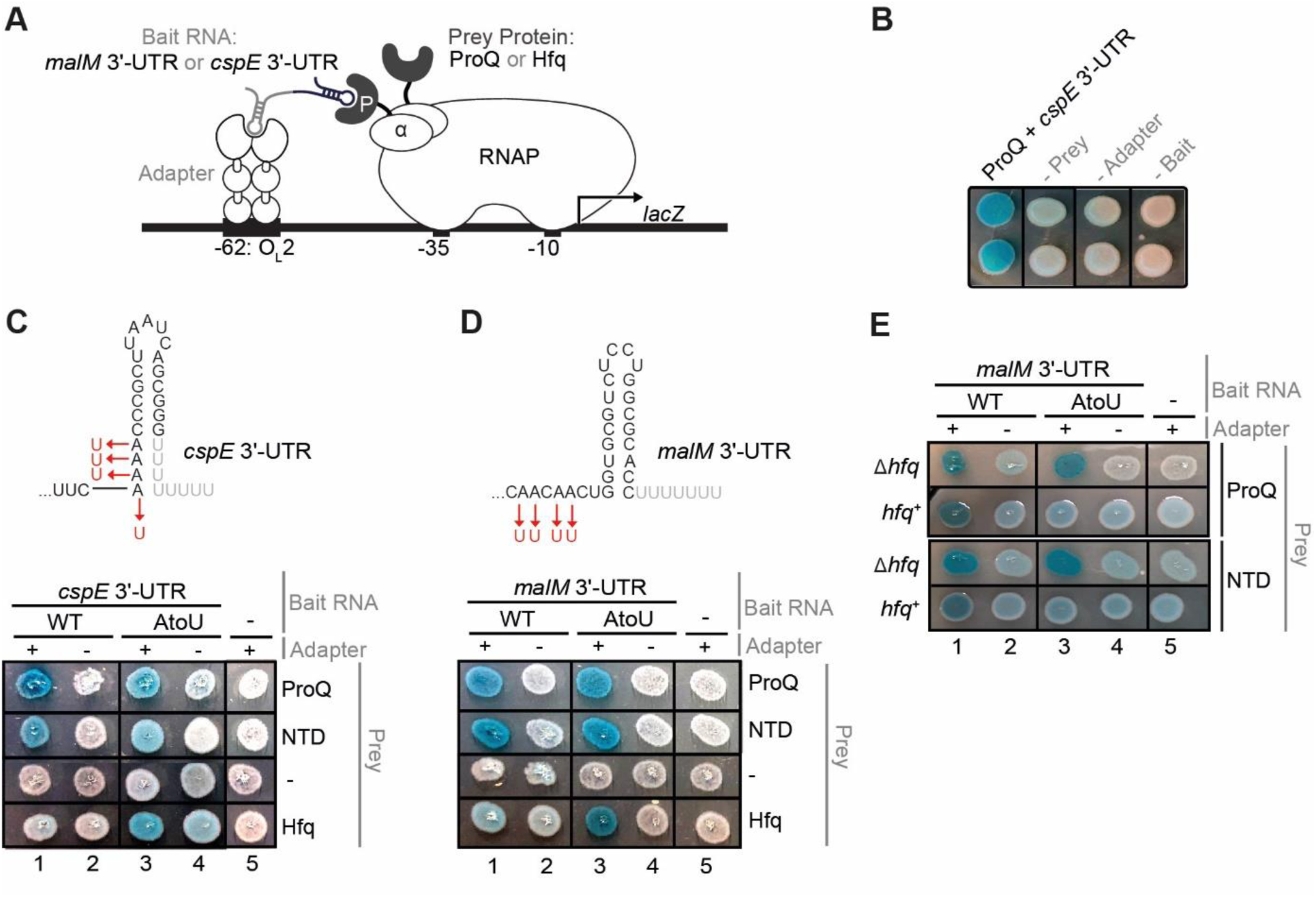
*In vivo* analysis of ProQ- and Hfq-binding determinants and cellular competition. (A) Design of a bacterial three-hybrid assay (B3H) system to detect interaction between a protein (P) of interest (here, ProQ or Hfq) and an RNA (here *cspE* 3’-UTR or *malM* 3’-UTR). Interaction between the protein moiety and RNA moiety fused, respectively, to the NTD of the alpha subunit of RNAP (α) and to one copy of the MS2 RNA hairpin (MS2^hp^) activates transcription from test promoter, which directs transcription of a *lacZ* reporter gene. Compatible plasmids direct the synthesis of the α-fusion protein (under the control of an IPTG-inducible promoter), the CI-MS2^CP^ adapter protein (under the control of a constitutive promoter; pCW17) and the hybrid RNA (under the control of an arabinose-inducible promoter). (B) Colony-color phenotypes of B3H interaction between ProQ and *cspE* on X-gal-indicator medium. *Δhfq* reporter strain cells (KB483) were freshly transformed with three compatible plasmids: one that encoded λCI alone or the λCI-MS2^CP^ fusion protein, another that encoded α alone or an α-ProQ fusion protein, and a third that encoded a 1XMS2^hp^-cspE hybrid RNA or only the MS2^hp^ moiety. The left-most column (*cspE* + ProQ) contains all 3 hybrid plasmids, and each addition column contains two hybrid plasmids and one negative control plasmid: α alone (-Prey), λCI alone (-Adapter) or MS2^hp^ alone (-Bait). Transformed cells were spotted on LB-agar medium containing 0.2% arabinose, 1.5 μM IPTG, 40 μg/mL X-gal and 75 μM TPEG and grown overnight at 37°C. Duplicate spots are shown. (C, D) Top: Schematic of WT *cspE* 3’-UTR (C) or *malM* 3’-UTR (D) constructs and four adenosines that were mutated to uridines in the A-to-U mutants. Bottom: Results of a representative B3H plate-based assay detecting interactions between full-length ProQ, the N-terminal FinO domain of ProQ or Hfq with *cspE* (C) or *malM* (D) RNA. β-gal assays were performed with *Δhfq* reporter strain cells containing three compatible plasmids: one (Adapter) that encoded λCI (-) or the λCIMS2^CP^(+) fusion protein, another (Prey) that encoded α alone (-), α-ProQ (ProQ), α-ProQ-NTD (NTD), or α-Hfq, and a third (Bait RNA) that encoded an MS2^hp^-*cspE* (C) or MS2^hp^-*malM* (D) hybrid RNA - either WT or AtoU mutant - or an RNA that contained only the MS2hp moiety (-). (E) Results of a representative B3H plate-based assay detecting interactions between ProQ and its NTD with *malM* RNA. β-gal assays were performed with *Δhfq* or *hfq*^+^ reporter-strain cells containing three compatible plasmids, as above.

We began our B3H analysis with a fragment of *cspE* that has been previously used in ProQ B3H assays, and with Δ*hfq* reporter cells, which have yielded optimal signal-to-noise for B3H interactions with both ProQ and Hfq (30,51). Patches of Δ*hfq* reporter cells grown on X-gal-indicator medium appeared much more blue when all three hybrid components were present than in each of the three negative-control conditions when half of each component was absent (- Adapter, - Prey or - Bait; Fig. 12B). This increase in blueness indicates an increase in *lacZ* expression, which represents an interaction between *cspE* RNA and ProQ. Under these assay conditions, wt *cspE* 3’-UTR displays minimal interaction with Hfq (Fig. 12C, column 1), but strong interaction with ProQ (full length ProQ or NTD only), as previously reported (30).

The *cspE* 3’-UTR contains four adenosines that are predicted to form base pairs with four of the eight uridines of the intrinsic terminator (Fig. 12C, top). These adenosines were mutated to uridines and the interaction of wt and mutant RNA to both ProQ and Hfq was assessed using the B3H assay. A-to-U substitutions led to a substantial increase in RNA interaction with Hfq, without strongly affecting the degree of ProQ interaction (either full length ProQ or the NTD alone; Fig. 12C, bottom, column 3). These results indicate that both wt and A-to-U *cspE* variants are expressed inside of cells and are capable of interacting in the B3H assay, but that Hfq interaction is substantially strengthened by the A-to-U substitution.

We next tested whether the B3H assay could detect interactions of *malM* 3’-UTR with Hfq and ProQ, and whether adenosines just upstream of its terminator would play an analogous role to those in *cspE* 3’-UTR. The same fragment of *malM* used in gel-shift experiments above was cloned into the MS2^hp^ RNA-bait construct. Indeed, when all three hybrid components were present (MS2^hp^-*malM*, α-ProQ, and CI-MS2^CP^ adapter) in reporter cells, bacteria plated on X-gal-indicator medium appear much more blue than negative controls where half of each hybrid component is left out (Fig. 12D, column 1), indicative of a ProQ-*malM* interaction. As with *cspE*, wt *malM* 3’-UTR interacts more strongly in this system with ProQ than with Hfq. When the four adenosines upstream of the *malM* terminator were mutated to uridines, *in vivo* binding with Hfq was strengthened without altering the apparent interaction with ProQ (Fig. 12D, column 3). This *in vivo* result mirrors *in vitro* binding results (Figs. 10, 11).

The above *in vivo* experiments were conducted in the absence of endogenous Hfq, but in a native cellular context, ProQ may need to compete with Hfq for RNA binding, given their partially overlapping RNA ligands (11-13). We wondered whether competition with Hfq could affect how ProQ binds to RNAs lacking the A-rich motif. To test if such competition could be observed in the B3H system the binding of wt and mutant *malM* 3’-UTR to ProQ in *Δhfq* reporter cells was compared with the same interactions in reporter cells that express endogenous Hfq (*hfq*^*+*^; Fig 12E). Unlike in *Δhfq* cells where the ProQ-*malM* interaction appears unaffected by the A-to-U substitution, this substitution leads to an apparent loss of ProQ binding in *hfq*^*+*^ cells (Fig 12E). In other words, the presence of endogenous Hfq appears to have little effect on the interaction of ProQ with wt *malM* 3’-UTR, but strongly inhibits interaction of A-to-U *malM* 3’-UTR (Fig. 12E). This competitive effect was similar with either the isolated NTD or ProQ suggesting that, even in the presence of the CTD and linker, competition with Hfq may drive RNA interactions with ProQ *in vivo*. Together these results are consistent with our model in which adenosines upstream of the terminator are a positive determinant for ProQ interaction by reducing competition with Hfq.

## DISCUSSION

### Recognition of intrinsic terminators by the ProQ NTD

The data presented here demonstrate that the FinO domain of *E. coli* ProQ specifically recognizes intrinsic transcription terminator structures within its RNA ligands (Figs. 4, 5). This observation is consistent with RIL-seq and CLIP-seq studies showing that enriched RNAs were dominated by mRNA 3’-UTRs and sRNAs (11,12). In further support, the homologous domain of *L. pneumophila* RocC, and the full-length F-like plasmid FinO bound the Rho-independent terminators of their specific RNA ligands tighter than their 5’ parts (23,52). On the other hand, the *N. meningitidis* minimal ProQ, which is composed only of the FinO domain, recognizes both the intrinsic terminators and the structured DUS regions (24,25). Hence, it is possible that the detailed structural properties of FinO-like domains from different proteins could determine their ability to specifically recognize Rho-independent transcription terminators.

Given that the N-terminal FinO domain of ProQ appears to specifically bind to terminator structures, an interesting and important question is the role of the linker and CTD. Our data show that binding of ProQ is much less dependent on the presence of the 3’ terminator structure than the isolated NTD alone, suggesting that either the linker or CTD may participate in non-terminator-specific RNA binding. Previous studies have also indicated that the linker and CTD of ProQ may have more non-specific contributions to the binding (19,30,53), and that these contributions may vary between RNAs. Studies using HDX RNA binding analysis and *in vivo* three-hybrid assay showed that these regions of ProQ make a larger contribution to the binding of RNAs SraB and SibB, which are longer than *cspE* 3’- UTR (19,30). The linker region, together with the NTD, was also required for mRNA-dependent ProQ binding to the ribosomes (53). Hence, it is possible that the linker and the CTD could form additional contacts with the same RNA molecules bound by the NTD or with other nearby RNAs. We hypothesize that such contacts could be involved in promoting RNA-RNA interactions, such as the RNA pairs reported for ProQ in *E. coli* and *S. enterica* (12,14,15).

Within a Rho-independent terminator, our study suggests that the lower part of the hairpin and 3’ oligoU tract are essential for the binding to the ProQ NTD (Figs. 6,7). The observation that a four-uridine tail is sufficient for *malM*-3’ binding to the NTD (Fig. 7) is consistent with a recent observation that the single-stranded 3’ tails of ProQ-specific RNAs were shorter than those of Hfq-specific RNAs (11). However, the number of 3’-terminal uridines required for efficient ProQ binding may differ at the primary sequence level between RNAs, given that their secondary structures involve different numbers of uridines contributing to base pairs within their terminator hairpin. It is possible that the shorter length of 3’ oligoU tails of ProQ-specific RNAs could be one of the features that distinguish RNA ligands of ProQ from those of Hfq. The importance of the bottom part of the terminator stem and adjacent single-stranded sequences has also been observed in the interactions of the homologous FinO protein with FinP RNA (32,52). It was reported that both FinO and ProQ bound FinP terminator hairpins with single stranded extensions (29,52), and that the binding of FinO protected the lower part of the FinP RNA terminator hairpin from RNase degradation (32). Hence, these data suggest that the recognition of the junction including the lower part of the terminator hairpin and the 3’ attached oligoU tail could be a general property of RNA binding proteins from the FinO-domain family.

A study of RNA binding by ProQ mutants in *E. coli* suggested a model of RNA/ProQ interactions that could explain the roles of the lower part of the terminator stem and the 3’ U- tract in binding to the ProQ NTD (30). In this genetic screen for mutants affecting RNA binding by ProQ several conserved residues on the concave side of the NTD as well as in a β- hairpin at the intersection between concave and convex surfaces were found to be essential for the binding of *cspE* 3’-UTR and SibB sRNA (30). It has been proposed that the residues on the concave side of the NTD form an electrostatic scaffold that matches the diameter of an A-type double helix, and that it is possible for the 3’ tail of an intrinsic terminator bound on the concave side to wrap around the edge of the FinO domain towards the convex side (30). Such a model is consistent with our observation that both the bottom part of the stem and the 3’ tail are required for the RNA binding by the NTD (Figs. 6, 7), and with data showing the involvement of both sides of the FinO domain of the homologous FinO protein in binding to the FinP RNA (31).

### A terminator adjacent A-rich motif serves as a negative determinant for Hfq binding

The strongest RNA ligands of ProQ both in *E. coli* and *S. enterica* datasets (9,11,12) have clear A-rich motifs on the 5’ side of the intrinsic terminator hairpin (Fig. 8A, C, Suppl. Fig. S7). This appears to be a distinctive feature of ProQ-specific RNAs, because in the strongest ligands of Hfq the same region is dominated by uridines (Fig. 8B) (9,12). While Hfq can bind A-rich sequences, for example ARN motifs, in some of its RNA ligands, these sequences are located either in mRNA 5’-UTRs (35-37) or far upstream of intrinsic terminators in a subgroup of Hfq-specific sRNAs, called Class II sRNAs (40-42). However, in the dominant subgroup of Hfq-binding sRNAs, called Class I sRNAs, the Hfq-binding module consists of a U-rich motif just upstream of the intrinsic terminator, which is followed by a long 3’ oligoU- tail (39,54). Hence, it appears that the presence of the A-rich motif immediately upstream of the transcription terminator is a feature that is unique to ProQ-specific RNAs relative to Hfq-specific RNAs.

Our data presented here support a model in which the A-rich motif upstream of the terminator hairpin serves as a negative determinant of binding to the Hfq protein (Figs. 9-12, Suppl. Fig. S7). One reason for the detrimental effect of this motif on Hfq binding may come from the fact that adenosines of the A-rich motif base-pair to uridines of the 3’ oligoU tail (Fig. 9). As a result, the 3’ oligoU tail becomes involved in secondary structure, which makes it less available for binding to Hfq. This agrees with the observation that single-stranded oligoU tails of ProQ-binding RNAs are shorter than those of Hfq-binding RNAs (11). On the other hand, the absence of an U-rich motif, on the 5’ side of the terminator in ProQ-specific RNAs, likely prevents interactions with the rim of Hfq, which are known to contribute to Hfq-RNA interactions (39,54). Hence, regardless of which of these two mechanisms is more important, the presence of the A-rich sequence on the 5’ side of the terminator would weaken the RNA binding to Hfq, in agreement with our *in vitro* and *in vivo* data (Figs. 11, 12). This suggests that correct recognition of RNA ligands by ProQ is dependent in part on competition with Hfq. Indeed, our *in vivo* results with both *cspE* and *malM* 3’-UTRs (Fig. 12) are consistent with adenosines immediately upstream of an intrinsic terminator being a positive determinant for ProQ binding – not through strengthening direct interactions with ProQ – but by serving as a negative determinant for Hfq binding, minimizing competition with this alternative RNA-binding protein.

In summary, we propose that recognition of RNAs by ProQ depends not only on the specific binding of the FinO-like NTD to select structures but also on competition with the Hfq protein. We show that the presence of the A-rich motif on the 5’ side of the intrinsic transcription terminators serves to increase their availability for binding to ProQ by limiting the access of Hfq. Further studies are needed to reveal how the functional differences between ProQ and Hfq correlate with which of them is preferred for binding at 3’ ends of individual RNAs. Interestingly, other proteins from this family, including FinO (21), FopA (J. Vogel, personal communication), meningococcal ProQ (25), and RocC (23), have different RNA binding specificities than ProQ (17). It remains to be seen if this RNA recognition specificity is determined by motifs in their RNA ligands, or rather by differences in the structure of FinO domains and their N- or C-terminal extensions.

## SUPPLEMENTAL DATA

Supplemental data are available as a PDF file on-line. These include raw gel images related to figures in the main text, supplemental data on the NTD binding to *cspE*-3’, the list of sequences used in RNA sequence comparisons, and the lists of oligonucleotides and bacterial strains used in the experiments.

## AUTHOR CONTRIBUTIONS

E.M.S. analyzed RNA binding by ProQ and NTD, and cloned and purified both proteins, J.K. analyzed RNA binding by Hfq, E.M.S., J.K., M.O. analyzed the binding data, M. M. B. analyzed the nucleotide frequency of A-rich sequences, C.M.G. and K.E.B analyzed RNA binding *in vivo* using bacterial three-hybrid assay, E.M.S., J.K., K.E.B., M.O. wrote the paper.

## ACKNOWLEDGEMENTS

We thank Gisela Storz for sharing unpublished data, helpful discussions, and critical comments on the manuscript. We thank Erik Holmqvist and Olke Uhlenbeck for helpful discussions. We thank Jakub Kosicki and Lechoslaw Kuczynski for advice on statistical analysis of RNA sequence comparisons, and Michal Szczesniak for advice on computational methods.

