## Supplemental data for "Determinants of RNA recognition by the FinO domain of the *Escherichia coli* ProQ protein"

### SUPPLEMENTAL MATERIALS AND METHODS

#### ProQ-binding assay

The RNA binding to ProQ and NTD proteins was measured using a gelshift assay as described in the Materials and Methods section in the main text.

#### Hfq-binding assay

The RNA binding to the Hfq protein was measured using a double-filter retention assay as described in the Materials and Methods section in the main text.

### SUPPLEMENTAL TABLES

**Supplemental Table S1. DNA oligonucleotides used to prepare templates for *in vitro* transcription.**

| **Name** | **Sequence (5’ → 3’)** |
|---|---|
| cspE-3′_F | TAATACGACTCACTATAGTAAGATACGTCAGCAAGAATTCAAAACCCGCTTAATC |
| cspE-3′_R | AAAAAAAACCCGCTGATTAAGCGGGTTTTGAATTCTTGCTGACGTATCTTAC |
| cspE-3′-ext+5_R | AGACCAAAAAAAACCCGCTGATTAAGCGGGTTTTGAATTC |
| cspE-3′-ext+22_R | CCTCAACCGCACACTAAAGACCAAAAAAAACCCGCTGATTAAGCGGGTTTTGAATTC |
| cspE-3′-6U_R | AAAAAACCCGCTGATTAAGCGGGTTTTGAATTCTTGCTGACGTATCTTAC |
| cspE-3′-4U_R | AAAACCCGCTGATTAAGCGGGTTTTGAATTCTTGCTGACGTATCTTAC |
| cspE-3′-0U_R | CCCGCTGATTAAGCGGGTTTTGAATTCTTGCTGACGTATCTTAC |
| cspE-3′-AtoU_F | TAATACGACTCACTATAGTAAGATACGTCAGCAAGAATTCTTTTCCCGCTTAATC |
| cspE-3′-AtoU_R | AAAAAAAACCCGCTGATTAAGCGGGAAAAGAATTCTTGCTGACGTATCTTAC |
| cspE81- 3′_F | TAATACGACTCACTATAGGCCCTTCTGCTGCAAACGTAATCGCTCTGTA |

| | |
|---|---|
| | AGATACGTCAGCAAG |
| cspE81-3′_R | AAAAAAAACCCGCTGATTAAGCGGGTTTTGAATTCTTGCTGACGTATCTTACAGAG |
| cspE81-3′-AtoU_R | AAAAAAAACCCGCTGATTAAGCGGGAAAAGAATTCTTGCTGACGTATCTTACAGAG |
| cspA-3′_F | TAATACGACTCACTATAGTAATCTCTGCTTAAAAGCACAGAATCTAAGATCCCTGCCATTTGGCGGGGATTTTTTT |
| cspA-3′_R | AAAAAAATCCCCGCCAAATGGCAGGGATCTTAGATTCTGTGCTTTTAAGCAGAGATTAC |
| cspA-3′-AtoR_F | TAATACGACTCACTATAGTAATCTCTGCTTAAAAGCACAGAATCTTTTTTCCCTGCCATTTGGCGGGGATTTTTTT |
| cspA-3′- AtoR_R | AAAAAAATCCCCGCCAAATGGCAGGGAAAAAAGATTCTGTGCTTTTAAGCAGAGATTAC |
| gapA-3′_F | TAATACGACTCACTATAGGACCTGATCGCTCACATCTCCAAATAAGTTGAGATGACACTGT |
| gapA-3′_R | AAAAAAAAGAGCGACCGAAGTCGCTCTTTTTAGATCACAGTGTCATCTCAACTTA |
| gapA-3′-AtoU-R | AAAAAAAAGAGCGACCGAAGTCGCTCAAAAAAGATCACAGTGTCATCTCAACTTA |
| hupA-5′_F | TAATACGACTCACTATAGCGAAAAAAAGTGGCTATCGGTGCGTGTATGCAGGAGAGTGCTATTCTGGCATTTCCGTCGCACTCGATGCTTAGCAAGCG |
| hupA-5′_R | TTCATAAGTTATCCTTACAATGTGTTTATCGCTTGCTAAGCATCGAGTGCG |
| hupA-5′_mut_R | AAAACGTAAGTTATCCTTACGATGTGTTTATCGCTTGCTAAGCATCGAGTGCG |
| hupA-5′_term_R | AAAAAAAGGTGCGCCAGGAGACGCACCATTCATAAGTTATCCTTACAATGTGTTTATCGCTTGCTAAGCATCGAGTGCG |
| hupA-5′_hp_R | GGTGCGCCAGGAGACGCACCATTCATAAGTTATCCTTACAATGTGTTTATCGCTTGCTAAGCATCGAGTGCG |
| hupB-3′_F | TAATACGACTCACTATAGGTAAACTAAGCGTTGTCCCCAGTGGGGATGTG |
| hupB-3′_R | AAAAAAGGCACATCAGTAGATGCGCCCTTGAACTTCGTCACATCCCCACTGGGGAC |
| hupB-3′-RtoU_R | AAAAAAGGCACATCAGTAGATGCGCCAAAGAACTTCGTCACATCCCCACTGGGGAC |
| ihfA-3′_F | TAATACGACTCACTATAGCTTCGCCCAAAGACGAGTAATCTGAT |
| ihfA-3′_R | AGAAAAAAGGCCGCAGAGCGGCCTTTTTAGTTAGATCAGATTACTCGTC |
| ihfA-3′-AtoU_R | AGAAAAAAGGCCGCAGAGCGGCCAAAAAAGTTAGATCAGATTACTCGTC |
| infA-3′_F | TAATACGACTCACTATAGTCGCTGATTGTTTTACCGCCTGATGGGCGAAGAGAAA |
| infA-3′_R | ATAAAAAGGCCGGTTAAACCGACCTTTTACTCGTTCTTTCTCTTCGCCCATC |
| infA-3′-AtoU_R | ATAAAAAGGCCGGTTAAACCGACCAAAAACTCGTTCTTTCTCTTCGCCCATC |
| lpp-5′_F | TAATACGACTCACTATAGCTACATGGAGATTAACTCAATCTAGAGGGTATTAATAATGAAAGCTACTAAACTG |
| lpp-5′_R | AGTAGAACCCAGGATTACCGCGCCCAGTACCAGTTTAGTAGCTTTCATTATTAATACCCTC |
| lpp-5′-mut_R | AAAAGGGCGCGGGATTACCGCGCCCAGTACCAGTTTAGTAGCTTTCATT |

| | |
|---|---|
| | ATTAATACCCTC |
| lpp-5′-term_R | AAAAAAAGGTGCGCCAGGAGACGCACCAGTAGAACCCAGGATTACCGCGCCCAGTACCAGTTTAGTAGCTTTCATTATTAAT |
| lpp-5′-hp_R | GGTGCGCCAGGAGACGCACCAGTAGAACCCAGGATTACCGCGCCCAGTACCAGTTTAGTAGCTTTCATTATTAAT |
| malM-3′_F | TAATACGACTCACTATAGCTTTATCAGCAGTGTAAAAGGCAAGGGGTAATTACGCCCCACAGTG |
| malM-3′_R | AAAAAAAGGTGCGCCAGGAGACGCACCAGTTGTTGCAAAATCAGCACTGTGGGGCGTAATTA |
| malM-3′-ext+5_R | AGCATAAAAAAAGGTGCGCCAGGAGACGCACCAGTTGTTGCAAAATCAGCACTGTGGGGCGTAATTA |
| malM-3′-ext+17_R | TCCCAGGAAGGAAGCATAAAAAAAGGTGCGCCAGGAGACGCACCAGTTGTTGCAAAATCAGCACTGTGGGGCGTAATTA |
| *malM*-3′-bulge_R | AAAAAAAGGTGCGCCAGGAGACGTTTCACCAGTTGTTGCAAAATCAGCACTGTGGGGCGTAATTA |
| *malM*-3′-AA_R | AAAAAAAGGTTTGCCAGGAGACTTACCAGTTGTTGCAAAATCAGCACTGTGGGGCGTAATTA |
| *malM*-3′-UU_R | AAAAAAAGGTAAGCCAGGAGACAAACCAGTTGTTGCAAAATCAGCACTGTGGGGCGTAATTA |
| *malM*-3′-4bp_R | AAAAAAAGGTGAGGACACCAGTTGTTGCAAAATCAGCACTGTGGGGCGTAATTA |
| *malM*-3′-3bp_R | AAAAAAAGGTAGGAACCAGTTGTTGCAAAATCAGCACTGTGGGGCGTAATTA |
| *malM*-3′-2bp_R | AAAAAAAGGAGGACCAGTTGTTGCAAAATCAGCACTGTGGGGCGTAATTA |
| *malM*-3′-1bp_R | AAAAAAAGAGGACAGTTGTTGCAAAATCAGCACTGTGGGGCGTAATTA |
| malM-3′-9U_R | AAAAAAAAAGGTGCGCCAGGAGACGCACCAGTTGTTGCAAAATCAGCACTGTGGGGCGTAATTA |
| malM-3′-6U_R | AAAAAAGGTGCGCCAGGAGACGCACCAGTTGTTGCAAAATCAGCACTGTGGGGCGTAATTA |
| malM-3′-5U_R | AAAAAGGTGCGCCAGGAGACGCACCAGTTGTTGCAAAATCAGCACTGTGGGGCGTAATTA |
| malM-3′-4U_R | AAAAGGTGCGCCAGGAGACGCACCAGTTGTTGCAAAATCAGCACTGTGGGGCGTAATTA |
| malM-3′-3U_R | AAAGGTGCGCCAGGAGACGCACCAGTTGTTGCAAAATCAGCACTGTGGGGCGTAATTA |
| malM-3′-2U_R | AAGGTGCGCCAGGAGACGCACCAGTTGTTGCAAAATCAGCACTGTGGGGCGTAATTA |
| malM-3′-1U_R | AGGTGCGCCAGGAGACGCACCAGTTGTTGCAAAATCAGCACTGTGGGGCGTAATTA |
| malM-3′-AtoU_R | AAAAAAAGGTGCGCCAGGAGACGCACCAGAAGAAGCAAAATCAGCACTGTGGGGCGTAATTA |
| micA_F | TAATACGACTCACTATAGAAAGACGCGCATTTGTTATCATCATCCCTGAATTC |
| micA_R | AAAAGGCCACTCGTGAGTGGCCAAAATTTCATCTCTGAATTCAGGGATGATGAT |
| sibA_F | TAATACGACTCACTATAGAGGGTTAGGGAGAGGTTTCCCCCTCCCCCTGGTGTTCTTAGTAAGCCTGGAAGCTAATCACTAAGAGTA |
| sibA_R | GGGAAAGCCTCTCCCGGAGAAGAGGGCTTTTAATAAGGAAAGGGTTATGATGAAGCACGTCATCATACTGGTGATACTCTTAGTGATTAGCT |

**Supplemental Table S2. DNA oligonucleotides used to prepare the insert to clone *E.coli proq* gene into the pET-15b expression plasmid.**

| **Name** | **Sequence (5’ → 3’)** | **Additional information** |
|---|---|---|
| ProQ/NTD_1F | GAAAATCTTTATTTCCAATCCATGGAAAATC | insertion of TEV Protease recognition site on the 5′ side of the protein coding sequence |
| ProQ_1R | **GGATCC**TCAGAACACCAGGTGTTCTGCGC | insertion of BamHI restriction site on the 3′ side of the protein coding sequence |
| NTD_1R | GTTAGCAGCC**GGATCC**TTACTCACCAG | |
| ProQ/NTD_2F | GCAGCCATATGCTCGA**GGATCC**gGAAAATCTTT | insertion of BamHI restriction site on the 5′ side of the protein coding sequence |
| ProQ_2R | GCAATACATCAC**GGATCC**TCAGAACACCAG | additional sequence on the 3′ side of the insert to ensure efficient BamHI cleavage |
| NTD_2R | GTTTAACGGGCTTTGTTAGCAGCC**GG** | |

GAAAATCTTTATTTCCAATCC – TEV Protease recognition sequence

**GGATCC** – BamHI restriction site

g – introduced to maintain the open reading frame

XXXXXX – sequences inserted to improve the efficiency of BamHI cleavage

XXXXXX – ProQ/NTD coding sequence

**Supplemental Table S3. Sequences on the 5' side of the terminators of ProQ- and Hfq-specific RNAs, which were used for the sequence logo analysis.**

| **Top 50 *E. coli* ProQ ligands RIL-seq dataset (1)** | **sequence** | **Top 50 *E. coli* Hfq ligands RIL-seq dataset (1)** | **sequence** | **Top 50 *E. coli* ProQ ligands CLIPseq dataset (2)** | **sequence** | **Top 50 *S. enterica* ProQ ligands CLIPseq dataset (2)** | **sequence** | **Top 50 *S. enterica* Hfq ligands CLIPseq dataset (3)** | **sequence** |
|---|---|---|---|---|---|---|---|---|---|
| malM 3' | UGCAACAACU | chiX | CUCUUUGACG | cspC 3' | UCGCUAUAAA | fliC 3' | ACACAAUCAA | chiX | CUCUUUGACG |
| acpP 3' | UGAACAUCUC | sroC | AACAAUCUUC | uspA 3' | UGUUCCUGAA | hupB 3' | AAAGUACAAG | invH 3'/invR | CUUUUUGAUA |
| ryfA | UAAGCCCGAA | flgL 3' | AUGUUUUGUC | csrB | GAAACGAACC | uspA 3' | UCAACAACAA | rprA | UGUAGUCUUU |
| cspE 3' | GAAUUCAAAA | omrA | CUUGCACCAA | gadC 3' | UAUAAGACAA | csrC | CAAGAAAAAA | oxyS | GUGAACUUUU |
| cspA 3' | GAAUCUAAGA | uhpT 3' | GGUGACUUUU | tnaA 3' | UAAUACUACA | ompD 3' | UGAAUACAAA | sdsR | CCAUUUCCCU |
| rybB | UUUUGUGGAG | rydC | CCGUAUUCUU | cspD 3' | AAAGGCAAAA | ompA 3' | UGAUAAAAAA | omrA | CUUGCACCAA |
| raiZ * | GUAUCGCCAA | glnA 3' | GCUAUCUGUA | sibC | UCUCCUUCAA | atpC 3' | AAGCAUAAAA | rybB | UUUUGUGGAG |
| hupB 3' | GAAGUUCAAG | rybB | UUUUGUGGAG | lpp 3' | AAGUGAAAAA | ecnB 3' | UAAUCAUCAA | sroC | CAAUCUUCGA |
| infA 3' | ACGAGUAAAA | sucD 3' | CGACAUGGUU | manZ 3' | UGUACACUAC | cspE 3' | GAUUUCAAAA | glmZ | CCUGUUUUGA |
| ihfA 3' | UAACUAAAAA | ykgH 3' | UGACAAUGAC | mcaS | GUGUACUGUA | csrB | GAAACGAACC | STnc2080 | CAUGACCCAU |
| cspD 3' | AAAGGCAAAA | arcZ | GUAUUCGCGC | cspE 3' | GGAUUCAAAA | pckA 3' | UGAGAAGAAA | arcZ | UAUUCGCGCA |
| garD 3' | CCUCGCAAAA | cyaR | CACCUCCUUA | rmf 3' | CUUUUAAAAA | manZ 3' | UGUACACUAC | mgrR | CAAGCAUUUA |
| lpp 3' | AAGUGAAAAA | gadY | CUAUCCUCUU | ompC 3' | AUCGAACAAA | lppA 3' | GUAAUAAAAA | micF | CCUAUUUCAA |
| manZ 3' | UGUACACUAC | micL | GCUCGUACCA | dps 3' | CCGAUAAAAG | mdh 3' | CCUGCCCGAU | STnc290 | ACAUGUUUAC |
| ompF 3' | GCCGAAAAAA | micF | CCCUAUUUCA | malM 3' | UGCAACAACU | acpP 3' | AAGUGAACAU | fnrS | UUCGUACGAC |
| fbaA 3' | CUUAUCUCAA | rprA | UGUAGUCUUU | upd 3' | CCUGUCUGAA | sodA | UCUGUAAGAA | spf | UAUUUUAGCC |
| rraB 3' | AAUGUUCGAA | dsrA | GCAAGUUUCA | rpsQ 3' | UACGAAUAAA | SL1344_0757 3' | UUUCCCCAAA | flgL 3'/STnc840 | AUGUUUUGUC |
| mcaS | GUGUACUGUA | mcaS | GUGUACUGUA | hupB 3' | GAAGUUCAAG | ryfD | AAAGAAUCAA | cpxP 3'/STnc870 | UCCCUGUCUU |
| adhE 3' | AGGCCUAUAA | sdsR | CAUUUCCCUG | talB 3' | UAAUCAUUCU | gltA 3' | UAAUUGACAA | cyaR | CACCUCCUUG |
| sodB 3' | GCAACUGUAA | gcvB | GAUUAAUGUA | csrC | CAGGAAAAAA | SL1344_1243 3' | UAAUUUAAGA | rydC | CGUCCUCCUU |
| clpA 3' | GGUUGGUCAA | malM 3' | UGCAACAACU | ompA 3' | GGUAGAAAAA | prs 3' | GGAUCUGAAA | ylbA 3' | GUUGGAUUUU |
| grcA 3' | AAGUUACUAA | dicF | GGUGACGUUU | acpP 3' | AAGUGAACAU | aspA 3' | GUACCAAAGA | sucD 3' | CGACAUGGUU |
| glpC 3' | UCCUUUCUGA | micA | AUGAAAUUUU | ecnB 3' | UAAGCAAUAA | rpoS 3' | GUCAAAAAAA | spf | AAUCGGAUUU |
| ynhF 3' | GAUUUCCUCU | omrB | AAUUACACCA | yqjK 3' | ACUUCCCUCU | lpxC | GUAUAAAAUU | micA | AUGAAAUUUU |

| | | | | | | | | | |
|---|---|---|---|---|---|---|---|---|---|
| cspC 3' | UCGCUAUAAA | cpxP | UCCCUGUCUU | rpsA 3' | CAAGCACUAA | fba 3' | CUGAUUGAAA | dsrA | GCAAGUUUGA |
| ldtE 3' | ACAAAAAAAA | lgoD 3' | AGUUGCCAUG | glgS 3' | ACUGAUAUAA | SL1344_4176 3' | CUUAUGAAUG | misC | AAAUUGACAA |
| icd 3' | AAUUUAUUAA | mgrR | CAAGCAUUCA | ahpC 3' | ACCUAAUUCU | rplQ 3' | CGUAAAAAAA | SL1344_1149 3' | CUGCAAGUUA |
| dapD 3' | AGUAUGCACA | sgrS | AAAUCACCCG | sokB | UUCAUUCGUU | glmS 3' | CCAGGCAGAA | dnaJ 3' | UUUUUCUUCA |
| mgaS 3' | UAAUAUUGCA | fnrS | UUCGUACGAC | ytjA 3' | UUUAUCCAAA | yejM 3' | CCUCGAAGAA | STnc600 | CUAUCUCCUU |
| acnB 3' | AACGACACAA | ytfK 3' | AGUAAUAAGA | rpoS 3' | CACAAUAAAA | tuf 3' | GAAUUGAAGA | SL1344_0036 3' | GCGAUGCCCA |
| ppa 3' | GGCGUAAUAA | spf | AUAUUUUAGC | sibB | CCUUAUUUAU | gstA 3' | AUGUCAUGUU | sgrS | AAAAUCACCC |
| flxA 3' | CAGCCGUAUU | malG 3' | ACUGUUAUUC | icdC 3' | AAUAUUGAAA | ompF 3' | GCCUCAAAAA | narK 3' | AUUAUCCUUU |
| bolA 3' | UUAUGAAAAA | oxyS | GUGAACUUUU | glmY | CCAUAACAAA | uspG 3' | AAACUUACAC | SL1344_0083 3' | AAUCCACUCA |
| pflB 3' | UCCCCGUAAA | ryhB | GUAUUACUUA | gapA 3' | GAUCUAAAAA | cheZ 3' | CGUCAACCCG | gcvB | UUCAGACCAC |
| uspE 3' | UAACUGACAA | tdcG 3' | UUCAUCCUUG | gcvB | GAUUAAUGUA | carB 3' | AAUGUUGAAA | rfrB | AGUAUUAUUU |
| yciB 3' | AAAUCCUAAC | dinI 3' | GAUUUUUUUU | dnaK 3' | AUUAUACUGA | sdsR | CAUUUCCCUG | ilvL | CGAACUAAGA |
| hns 3' | UUGCUUAAAA | yehA 3' | GUCUGCAUUU | spy 3' | AAACUUAAGA | int 3' | UAGAAAAGAA | adiA 3' | AUACCUUUUU |
| yfcZ 3' | CGGGUACGUA | allE 3' | GUUGGAUUUU | ihfA 3' | UAACUAAAAA | rpsI 3' | AGCGAAAAAA | ybiJ 3' | ACCCUUAUUU |
| uhpT 3' | GGUGACUUUU | G0-10699 | AAAUGGCGUC | lpxC 3' | GUAUAAAAUU | osmE 3' | UUCAAGGAAA | ryeF | GCUCGUACCA |
| cheW 3' | UUGAAAUGAA | cof 3' | CCUUCCAGCA | arrS | UUCGACUUAA | recA 3' | UUACCAUCAA | nlpB 3' | GUAAUUUAACA |
| atpC 3' | AAGCACAAAA | lpp 3' | AAGUGAAAAA | tdh 3' | ACACGAACAA | rpoH 3' | UGCCAAUGAA | gndA 3' | AAAUGUUUAA |
| cyaR | CACCUCCUUA | ygaM 3' | GCGUCUUUGA | frr 3' | ACGACAAAAA | stpA 3' | UAAAAACAUA | DapZ | UGAUUACAAA |
| mog 3' | GAAUAAAAAA | ryeG | UAUAGCUUGU | ytfL | AGUAAUAAGA | nusB 3' | GUGAUUCACA | arcZ | GUAUUCGCGC |
| rpoD 3' | UCUGCACAAA | sucB 3' | GCACUGUAGA | maoP 3' | UUAAUAAAAA | ssb 3' | UCUGCUGAAA | Cof | CUUCCAGCAA |
| ppnP 3' | GUAAUUCCUC | smpB 3' | AGUUCCUCAC | fbaA 3' | CUUAUCUCAA | lpdA 3' | AUACAAAAAA | ibpB | UCCUUGCCUU |
| gcvP 3' | AUUUUCUAAA | gadE | AGAACCCUUC | ptsG 3' | GGGAGACUAA | ihfA 3' | UAACUAAAAA | OmrB | AUUACACCAA |
| uspA 3' | UGUUCCUGAA | sroD | AGUCGUCAGA | eno 3' | GAUUUAAAAA | clpA 3' | GUCGUACAAA | osmE 3' | UUCAAGGAAA |
| tufA 3' | GAAUUGAAAA | rsmG 3' | UACAUUUAAC | aceA 3' | CUGACUGUAG | SL1344_3447 3' | UACAAAAAAA | STnc780 | CGUUUUAUAG |
| ompC 3' | AUCGAACAAA | ahpF 3' | CAAUUGCUUA | hchA 3' | CAUAAUCAAU | STnc1080 | GCACCAGAAA | STnc740 | GGCUGUAUUU |
| exuT 3' | UCCCUUCAAA | acpP 3' | UGAACAUCUC | rpmE 3' | UCCGAAAAAA | lon 3' | AUAAAAACAG | SL1344_1101 3' | UGAUUUCGGA |

**Supplemental Table S4. Bacterial strains used in bacterial three-hybrid assay.**

| Name | Relevant Details | Antibiotic | Source | Fig. |
|---|---|---|---|---|
| NEB5α F'Iq | *E. coli lacIq* host strain for plasmid construction | TetR | NEB | N/A |
| KB480 | FW102 harboring an F' episome bearing tetracycline resistance and test promoter *plac*-$O_L$2-62 fused to *lacZ* | TetR, StrR | (4) | Fig 12E |
| KB483 | FW102 *hfq::kan* harboring F' episome bearing tetracycline resistance and test promoter *lac*-$O_L$2-62 fused to *lacZ* | TetR, KanR, StrR | (4) | Fig 12 |

**Supplemental Table S5. Plasmids used in the bacterial three-hybrid assay**. Listed in alphabetical order by plasmid name.

| Name | Description | Details | Reference/ Source | Figures |
|---|---|---|---|---|
| pACλCI | empty vector | Encodes full-length λCI under the control of the *lacUV5* promoter; confers CamR | (5) | Fig 12 |
| pBrα | empty vector | Encodes full-length alpha under the control of tandem *lpp* and *lacUV5* promoters; confers AmpR | (5) | Fig 12 |
| pCH1 | empty vector (pCDF-1XMS2$^{hp}$) | Encodes a single MS2 hairpin (MS2$^{hp}$) in front of XmaI and HindIII restriction sites, under the control of the pBAD promoter; confers SpcR | (4) | Fig 12 |
| pCW17 | pAC-p$_{cons}$-λCIMS2$^{CP}$ | Encodes residues 1-248 of CI fused via a 12aa-linker to a MS2 coat protein (MS2$^{CP}$); transcription of fusion protein driven by a constitutive promoter; confers CamR | (4) | Fig 12 |
| pKB817 | pBrα-Hfq | Encodes residues 1-248 of alpha fused via three alanine residues to full-length wild-type *E. coli* Hfq; confers AmpR | (6) | Fig 12 |
| pKB949 | pBrα-ProQ$^{FL}$ | Encodes residues 1-248 of alpha fused via three alanine residues to full-length wild-type *E. coli proQ*; confers AmpR | (4) | Fig 12 |
| pKB1210 | pCDF-1XMS2$^{hp}$-*malM* | 3'UTR of *E. coli malM* (final 90 nts) cloned behind MS2$^{hp}$ in pCH1 between XmaI/ HindIII sites; RNA encodes its own terminator; confers SpcR | This study (oKB1532, oKB1533) | Fig 12A,D,E |
| pKB1257 | pCDF-1XMS2$^{hp}$-*malM* 4A::U | *malM* AACAA-->TTCTT mutation (see Table S6) introduced to pKB1210; confers SpcR | This study (oKB1580 + oKB1581) | Fig 12D,E |
| pKB1259 | pCDF-1XMS2$^{hp}$-*cspE* 4A::U | *cspE* AAAA → TTTT mutation (see Table S6) introduced to pSP10; confers SpcR | This study (oKB1584, oKB1585) | Fig 12C |
| pSP90 | pBrα-ProQ$^{NTD}$ | Encodes residues 1-248 of alpha fused via three alanine residues to residues 1-119 of wild-type *E. coli proQ;* TAA stop codon after proQ added through PCR; confers AmpR | (4) | Fig 12 |
| pSP10 | pCDF-1XMS2$^{hp}$-*cspE* | 3'UTR of *E. coli cspE* (final 85 nts) cloned behind MS2$^{hp}$ in pCH1 between XmaI/ HindIII sites; RNA encodes its own terminator; confers SpcR | (4) | Fig 12B,C |

**Supplemental Table S6. DNA oligonucleotides used in construction of plasmids for B3H assays.**

| Name | Description | Used for | Sequence |
|---|---|---|---|
| oKB1532 | F XmaI *malM* | PCR: pKB1210 | GGCCGGCCCGGGCTTTATCAGCAGTGTAAAAGGCAAGGG |
| oKB1533 | R HindIII *malM* | PCR: pKB1210 | CCGGCCAAGCTTAAAAAAAGGTGCGCCAGGAGACGC |
| oKB1580 | F *malM* A::U | Mut. PCR: pKB1257 | CTGATTTTGCTTCTTCTGGTGCGTCTCCTG |
| oKB1581 | R *malM* A::U | Mut. PCR: pKB1257 | CACTGTGGGGCGTAATTA |
| oKB1584 | F *cspE* A::U | Mut. PCR: pKB1259 | GCAAGAATTCTTTTCCCGCTTAATCAG |
| oKB1585 | R *cspE* A::U | Mut. PCR: pKB1259 | TGACGTATCTTACAGAGC |

**Supplemental Table S7. Sequences of hybrid RNAs used in B3H assay.** Light blue letters indicate linker and cloning sites. Dark blue letters indicate the 21-nt MS2$^{hp}$ sequence. Black letters indicate the bait RNA sequence inserted between XmaI and HindIII sites. Red letters indicate site-directed mutations introduced upstream of terminator hairpins.

| **Plasmid** | **Description** | **Sequence** |
|---|---|---|
| pSP10 | pCDF-1XMS2$^{hp}$-*cspE* 3'UTR | AGAAAACAUGAGGAUCACCCAUGUCUGCAGCCCGGGC AAAGGCCCUUCUGCUGCAAACGUAAUCGCUCUGUAAG AUACGUCAGCAAGAAUUCAAAACCCGCUUAAUCAGCG GGUUUUUUUU |
| pKB1210 | pCDF-1XMS2$^{hp}$-*malM* 3'UTR | AGAAAACAUGAGGAUCACCCAUGUCUGCAGCCCGGGC UUUAUCAGCAGUGUAAAAGGCAAGGGGUAAUUACGCC CCACAGUGCUGAUUUUGCAACAACUGGUGCGUCUCCU GGCGCACCUUUUUUU |
| pKB1257 | pCDF-1XMS2$^{hp}$-*malM* 4A::U | AGAAAACAUGAGGAUCACCCAUGUCUGCAGCCCGGGC UUUAUCAGCAGUGUAAAAGGCAAGGGGUAAUUACGCC CCACAGUGCUGAUUUUGC**UU**C**UU**CUGGUGCGUCUCCU GGCGCACCUUUUUUU |
| pKB1259 | pCDF-1XMS2$^{hp}$-*cspE* 4A::U | AGAAAACAUGAGGAUCACCCAUGUCUGCAGCCCGGGC AAAGGCCCUUCUGCUGCAAACGUAAUCGCUCUGUAAG AUACGUCAGCAAGAAUUC**UUUU**CCCGCUUAAUCAGCG GGUUUUUUUU |

**SUPPLEMENTAL FIGURES**

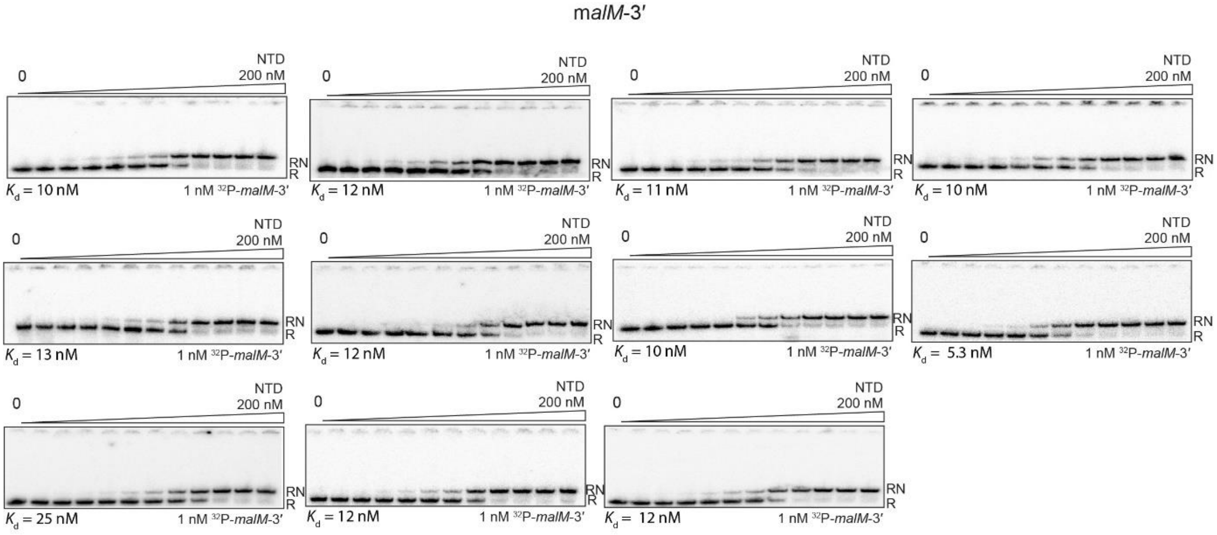


**Supplemental Figure S1. Gelshift analysis of the binding of 1 nM $^{32}$P-labeled *malM*-3′ to the N-terminal FinO domain of ProQ (NTD)**. The binding of wt *malM*-3′ to the N-terminal domain of ProQ was re-measured together with each batch of *malM*-3′ mutants to ensure that the affinity in the control experiment remains consistent. All individual experiments are shown. The fitting of the data from each experiment using the quadratic equation provided the $K_d$ values presented below the gels, while the average $K_d$ value is presented in the Table 1 in the main text. The concentration series of the NTD were made by 2-fold sequential dilutions. The used range of the NTD concentrations is indicated above the gels. Free $^{32}$P-RNA is marked as R, and RNA-NTD complex as RN.

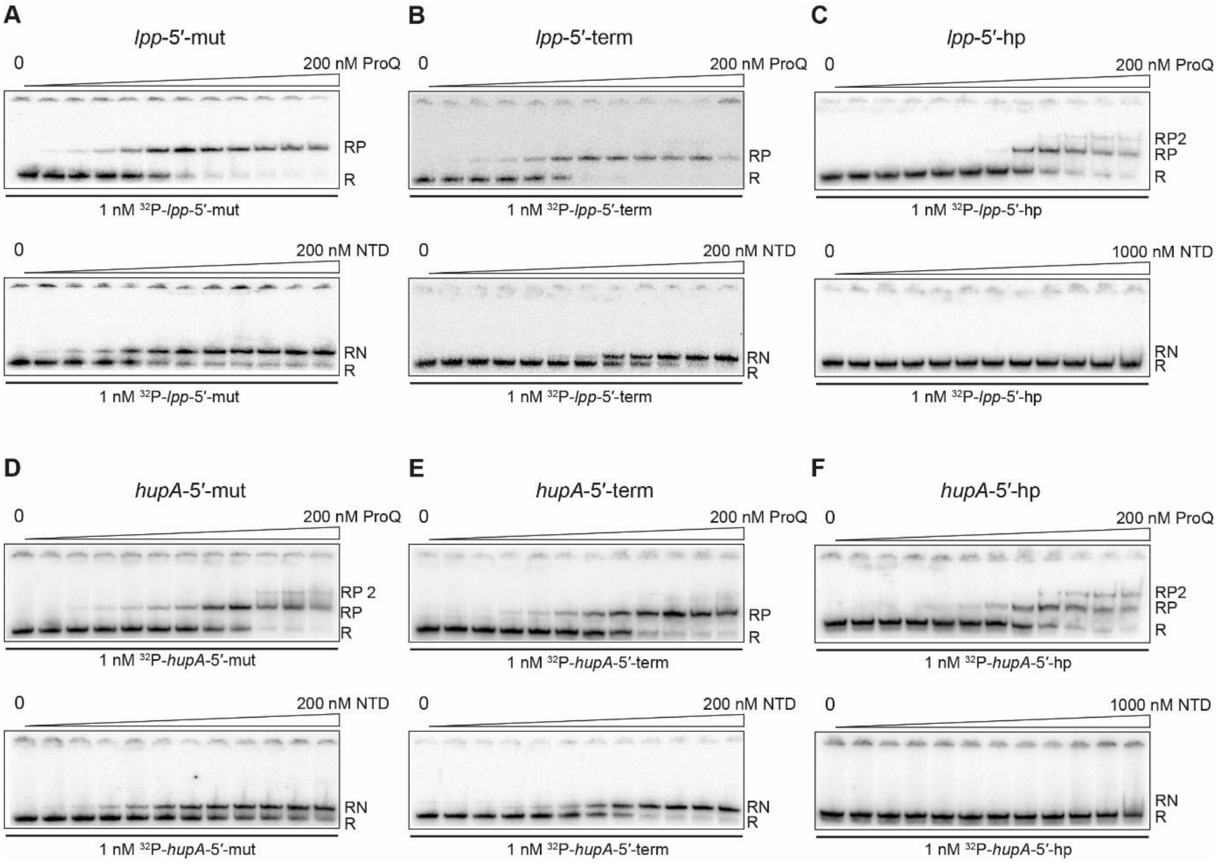


**Supplemental Figure S2. Gelshift analysis of the binding of *lpp*-5′-mut (A), *lpp*-5′-term (B), *lpp*-5′-hp (C), *hupA*-5′-mut (D), *hupA*-5′-term (E), and *hupA*-5′-hp (F) RNAs to the full-length ProQ protein or the ProQ NTD.** The raw data in the gels correspond to the data presented in plots on Fig. 4 in the main text. The concentration series of ProQ and NTD proteins were made by 2-fold sequential dilutions. The used ranges of the ProQ and NTD concentrations are indicated above the gels. Free $^{32}$P-RNA is marked as R, RNA-ProQ complex as RP, higher order RNA-ProQ complex as RP 2, and RNA-NTD complex as RN.

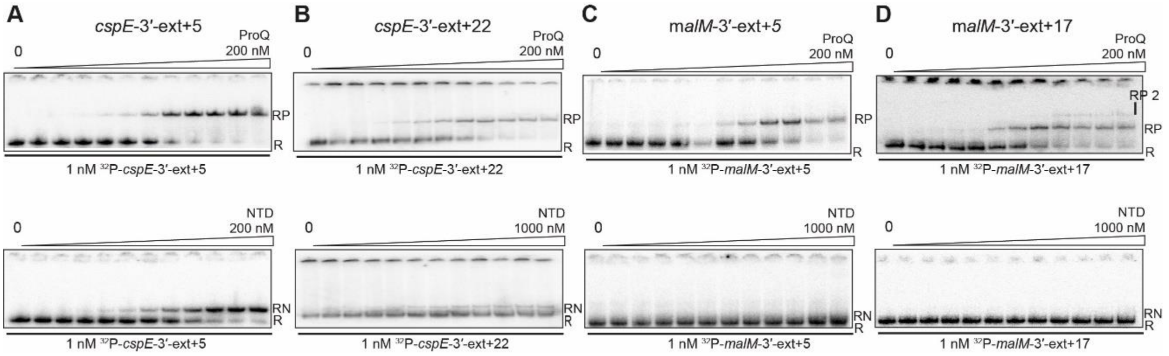


**Supplemental Figure S3. Gelshift analysis of the binding of *cspE*-3′-ext+5 (A), *cspE*-3′-ext+22 (B), *malM*-3′-ext+5 (C), and *malM*-3′-ext+17 (D) RNAs to the full-length ProQ protein or the ProQ NTD.** Raw data in the gels correspond to the data on the plot in Figure 5 in the main text. Descriptions are the same as in the Supplemental Figure S1.

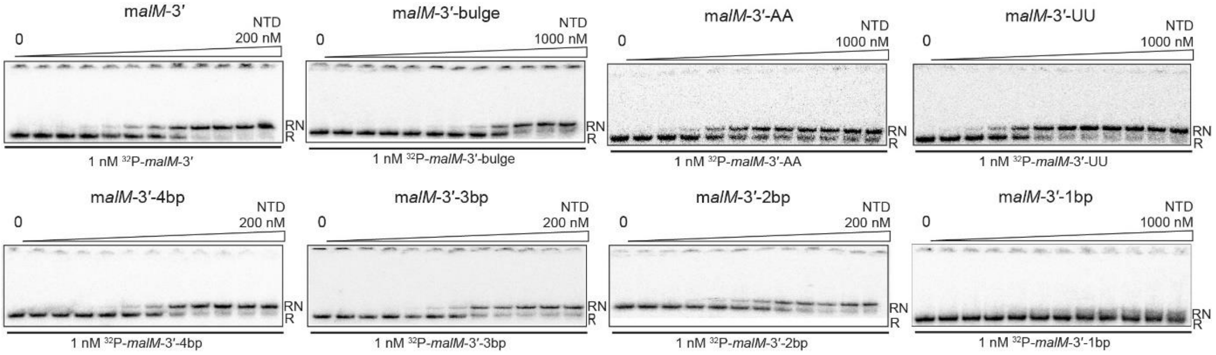


**Supplemental Figure S4. Gelshift analysis of the binding of *malM*-3′, *malM*-3′-bulge, *malM*-3′-AA, *malM*-3′-UU, *malM*-3′-4bp, *malM*-3′-3bp, *malM*-3′-2bp, and *malM*-3′-1bp to the ProQ NTD**. Raw data in the gels correspond to the data on the plot in Figure 6 in the main text. Descriptions are the same as in the Supplemental Figure S1.

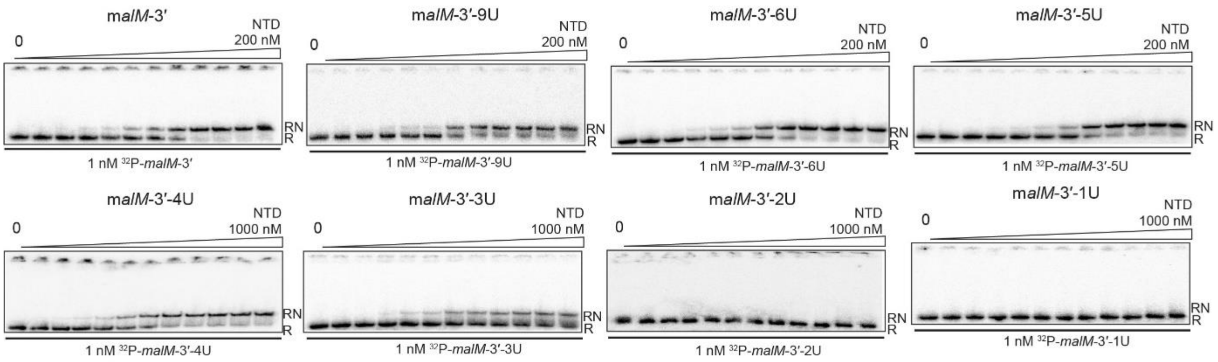


**Supplemental Figure S5. Gelshift analysis of the binding of *malM*-3′, *malM*-3′-9U, *malM*-3′-6U, *malM*-3′-5U, *malM*-3′-4U, *malM*-3′-3U, *malM*-3′-2U, and *malM*-3′-1U to the ProQ NTD**. The gels correspond to the data in the plot in Fig. 7 in the main text. Descriptions are the same as in the Supplemental Figure S1.

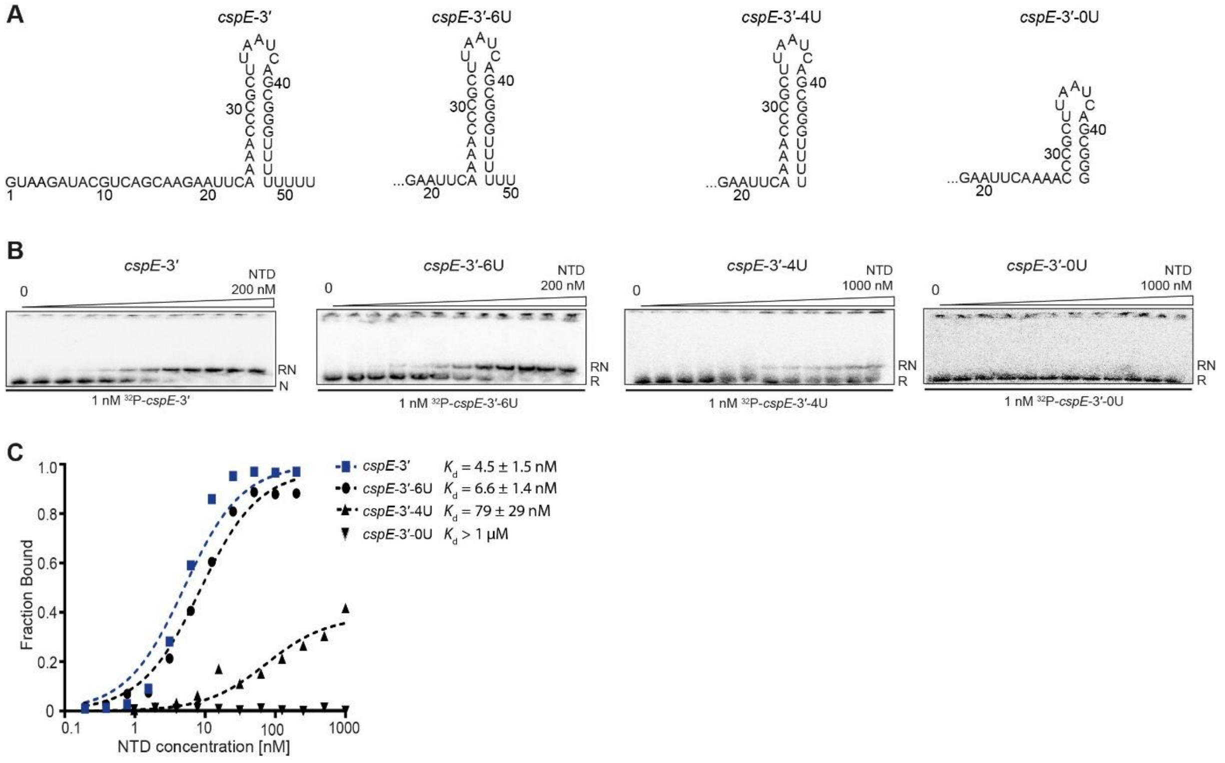


**Supplemental Figure S6. The effect of the length of 3′-terminal oligoU sequence on the binding of *cspE*-3′ to the ProQ NTD.** (A) The *cspE*-3′ constructs with shortened 3′-terminal oligouridine tails. (B) Equilibrium binding of 1 nM $^{32}$P-*cspE*-3′, $^{32}$P-*cspE*-3′-6U, $^{32}$P-*cspE*-3′-4U and $^{32}$P-*cspE*-3′-0U to the NTD monitored using gelshift assay. The NTD concentration series was prepared by 2-fold sequential dilutions, and the concentration range is indicated above the gels. Descriptions are the same as in the Supplemental Figure S1. (C) The fitting of the data from (B). Data for *cspE*-3′ are the same as in Figure 3 C (main text). The fitting of the data in the plot using the quadratic equation provided $K_d$ values of 7.9 nM *cspE*-3′-6U and 73 nM for *cspE*-3′-4U. The binding of *cspE*-3′-0U was essentially undetectable up to 1 µM concentration of the NTD. Average equilibrium dissociation constant ($K_d$) values with standard deviation from at least three independent experiments are shown in the legend.

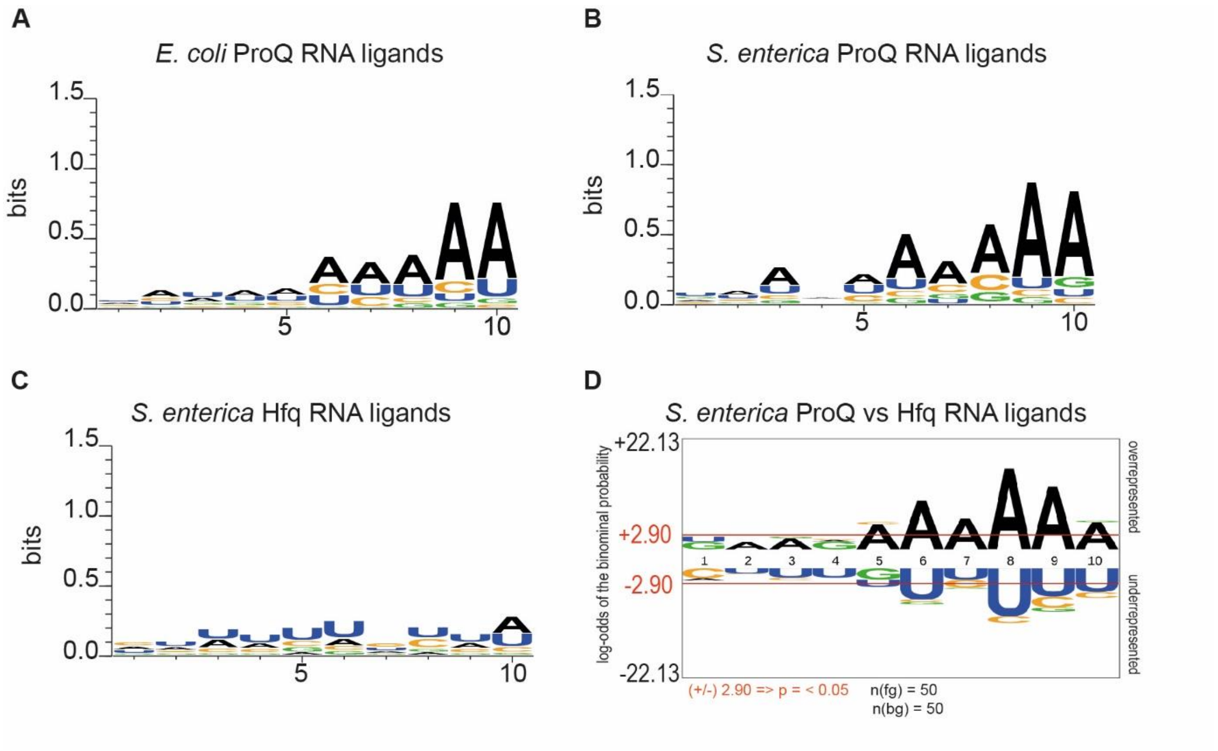


**Supplemental Fig. S7. Nucleotide frequency in the 10-nt sequence on the 5′ side of the terminator hairpin in RNA ligands of ProQ and Hfq proteins in previously reported *S. enterica* and *E. coli* datasets obtained using CLIP-seq (2,3).** (A) Nucleotide frequencies obtained by *WebLogo* for the 10-nt long sequence 5′-adjacent to the terminator hairpin of the top 50 RNA ligands of *E. coli* ProQ containing Rho-independent terminators, $p < 2.2 \times 10^{-16}$ (2); (B) Nucleotide frequencies obtained by *WebLogo* for the top 50 RNA ligands of *S. enterica* ProQ containing Rho-independent terminators, $p < 2.2 \times 10^{-16}$ (2); (C) Nucleotide frequencies obtained by *WebLogo* for the top 50 RNA ligands of *S. enterica* Hfq containing Rho-independent terminators, $p = 4.98 \times 10^{-16}$ (3); (D) The statistically significant nucleotide frequencies at each nucleotide positions for *S. enterica* ProQ RNA ligands as compared to Hfq RNA ligands analyzed by *pLogo* software for top 50 ProQ RNA ligands as a foreground, and top 50 Hfq RNA ligands as a background. Statistically significant p-values: A at position 5, $p = 4.15 \times 10^{-4}$; A at position 6, $p = 8.82 \times 10^{-9}$; A at position 7, $p = 3.24 \times 10^{-5}$; A at position 8, $p = 3.32 \times 10^{-15}$; A at position 9, $p = 1.24 \times 10^{-11}$; A at position 10, $p = 1.7 \times 10^{-4}$. For the selection criteria see Materials and Methods in the main text. The number of foreground sequences is marked on the figure as n(fg), and the number of background sequences as n(bg).

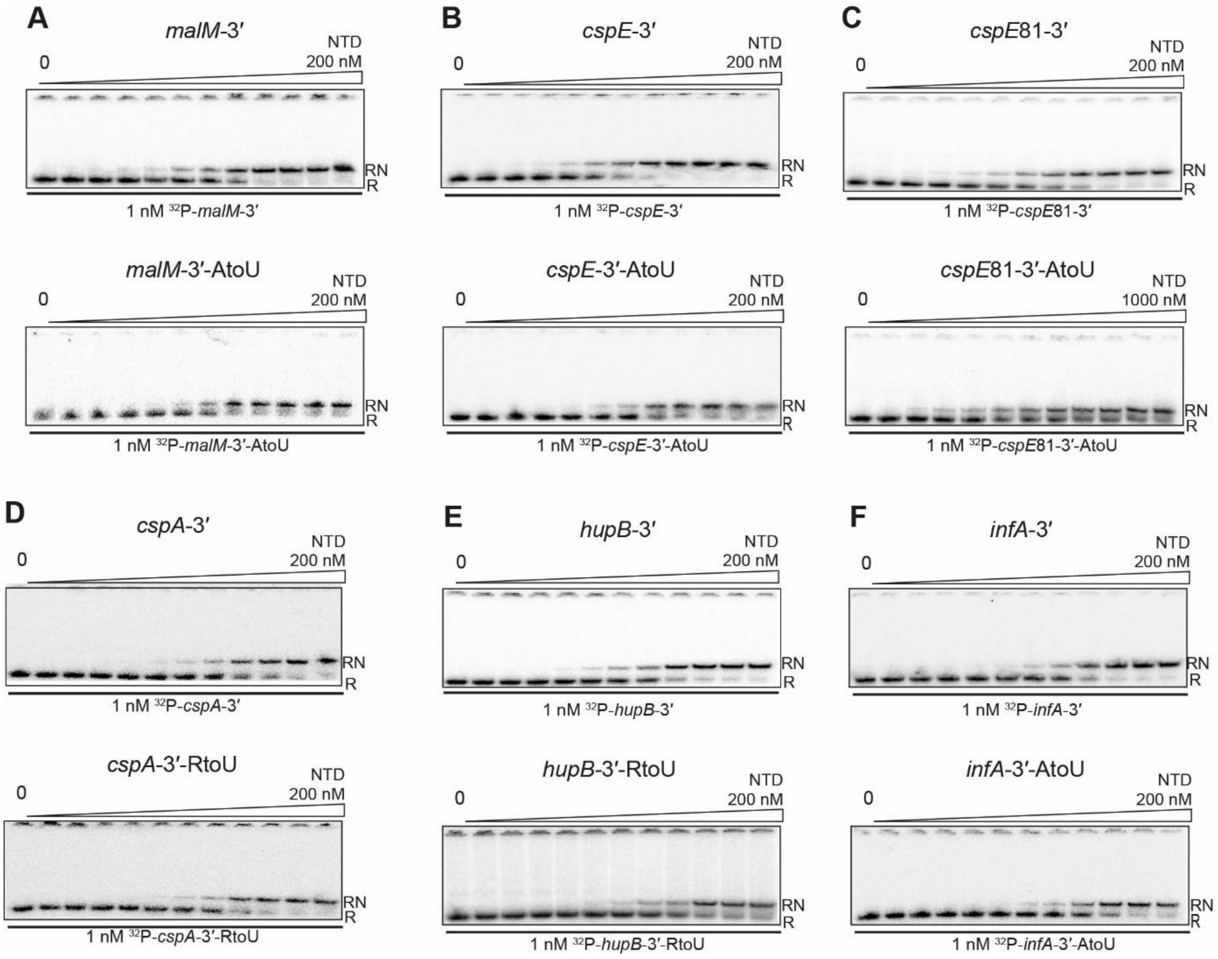


**Supplemental Figure S8. The binding of RNAs with mutations in the A-rich sequences to the ProQ NTD**. Gelshift analysis of the binding of *malM*-3′ and *malM*-3′-AtoU (A), *cspE*-3′ and *cspE*-3′-AtoU (B), *cspE*81-3′- and *cspE*81-3′-AtoU (C), *cspA*-3′ and *cspA*-3′-RtoU (D), *hupB*-3′ and *hupB*-3′-RtoU (E), *infA*-3′- and *infA*-3′-AtoU (F) RNAs to the ProQ NTD. The gels correspond to the data in the plots in Fig. 10 in the main text. Descriptions are the same as in the Supplemental Figure S1.

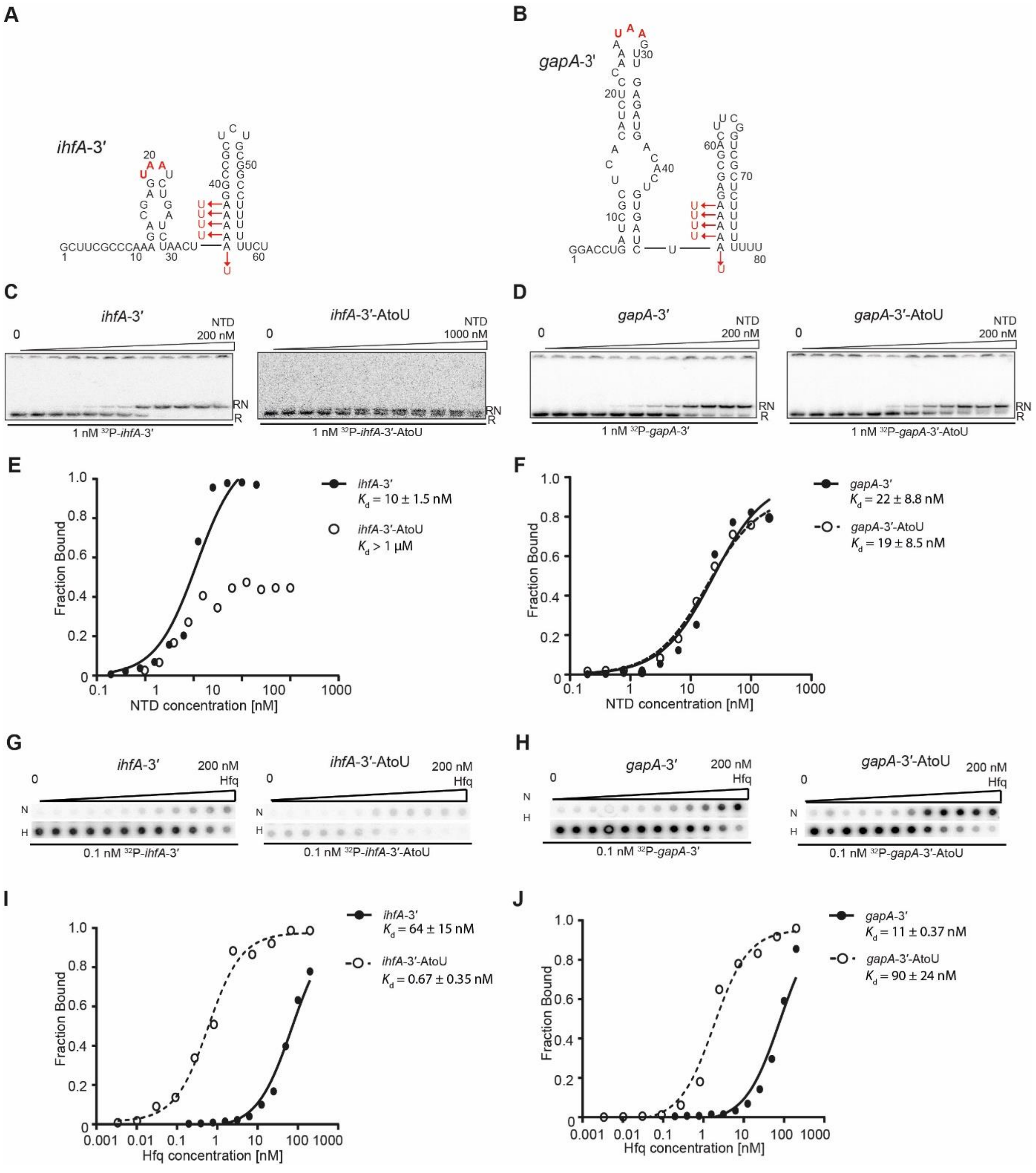


**Supplemental Figure S9. The binding of *ihfA*-3′ and *gapA*-3′ RNAs with mutations in the A-rich sequences to the ProQ NTD and to the Hfq protein**. In the panels A and B are shown secondary structures of *ihfA*-3′ (A) and *gapA*-3′ (B) RNAs with mutations introduced into the A-rich regions marked with red font. The data corresponding to the ProQ NTD binding monitored by gelshift assay are shown on panels C-F. This includes the gelshift analysis of the NTD binding of RNAs *ihfA*-3′ and *ihfA*-3′-AtoU (C) and *gapA*-3′ and *gapA*-3′-AtoU (D). The fitting of the data from C and D using the quadratic equation provided $K_d$ value of 11 nM for *ihfA*-3′, while the binding of *ihfA* -3′-AtoU was essentially undetectable up to 1 μM concentration of the NTD (E), and $K_d$ values of 23 nM for *gapA*-3′ and 18 nM for *gapA*-3′-AtoU (F). Descriptions are the same as in the Supplemental Figure S1. The data corresponding to the Hfq protein binding monitored by double-filter retention assay are shown on panels G-J. This includes the double-filter retention data of Hfq binding of RNAs *ihfA*-3′ and *ihfA*-3′-AtoU (G), and *gapA*-3′ and *gapA*-3′-AtoU (H). The fitting of the data from G and H using the quadratic equation provided $K_d$ values of 71 nM for *ihfA*-3′ and 0.55 nM for *ihfA*-3′-AtoU (I) and 79 nM for *gapA*-3′ and 1.8 nM for *gapA*-3′-AtoU (J). Average equilibrium dissociation constant ($K_d$) values from at least three independent experiments are shown in the legend.

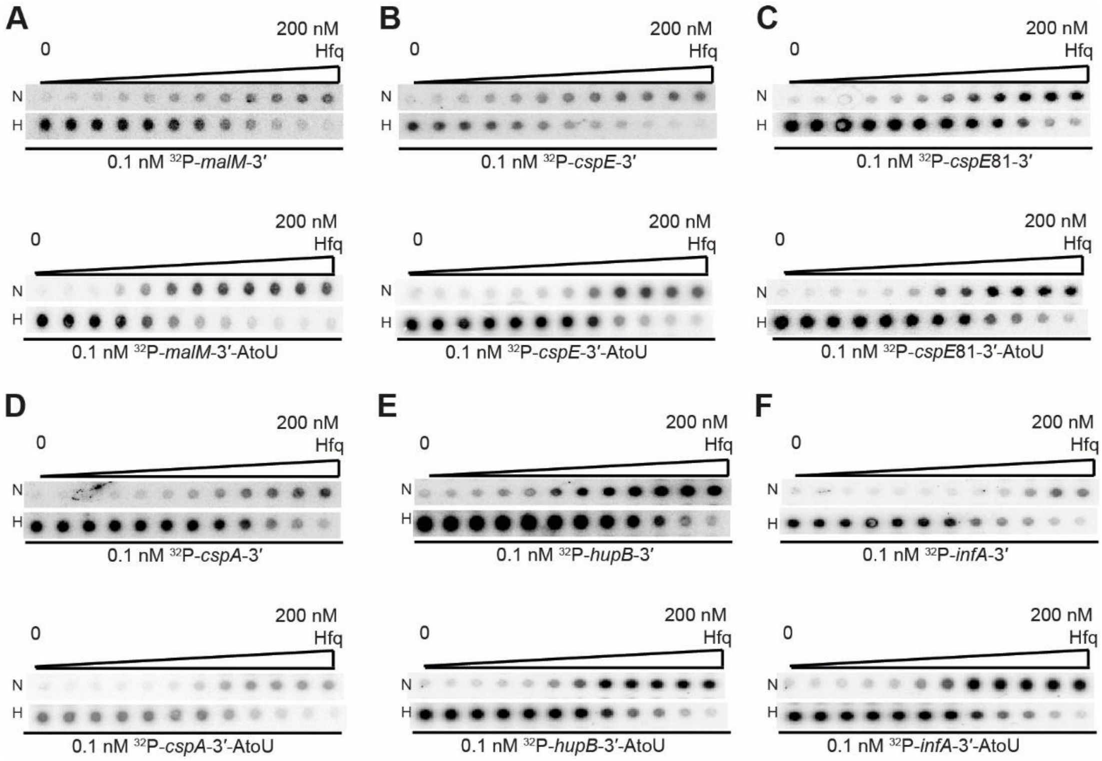


**Supplemental Figure S10. The binding of RNAs with mutations in the A-rich sequences to the Hfq protein**. The binding of $^{32}$P-labeled *malM*-3′ and *malM*-3′-AtoU (A), *cspE*-3′ and *cspE*-3′-AtoU (B), *cspE*81-3′ and *cspE*81-3′-AtoU (C), *cspA*-3′ and *cspA*-3′-AtoU (D), *hupB*-3′ and *hupB*-3′-AtoU (E), *infA*-3′ and *infA*-3′-AtoU (F) RNAs to Hfq protein using double-filter retention assay. Symbols N and H correspond to nitrocellulose and Hybond N+ membranes, respectively. The raw data in the double membranes correspond to the data presented in plots on Fig. 11 in the main text. The concentration series of Hfq protein were made by 2-fold sequential dilutions. The used ranges of the Hfq concentrations are indicated above the membranes.
